# Immunocapture of dsRNA-bound proteins provides insight into tobacco rattle virus replication complexes and reveals Arabidopsis DRB2 to be a wide-spectrum antiviral effector

**DOI:** 10.1101/842666

**Authors:** Marco Incarbone, Marion Clavel, Baptiste Monsion, Lauriane Kuhn, Helene Scheer, Vianney Poignavent, Patrice Dunoyer, Pascal Genschik, Christophe Ritzenthaler

## Abstract

Plant RNA viruses form highly organized membrane-bound virus replication complexes (VRCs) to replicate their genome and multiply. This process requires both virus- and host-encoded proteins and leads to the production of double-stranded RNA (dsRNA) intermediates of replication that trigger potent antiviral defenses in all eukaryotes. In this work, we describe the use of *A. thaliana* constitutively expressing GFP-tagged dsRNA-binding protein (B2:GFP) to pull down viral replicating RNA and associated proteins *in planta* upon infection with tobacco rattle virus (TRV). Mass spectrometry analysis of the dsRNA-B2:GFP-bound proteins from TRV-infected plants revealed the presence of (i) viral proteins such as the replicase, which attested to the successful isolation of VRCs, and (ii) a number of host proteins, some of which have previously been involved in virus infection. Among a set of nine selected such host candidate proteins, eight showed dramatic re-localization upon infection, and seven of these co-localized with B2-labeled TRV replication complexes, providing ample validation for the immunoprecipitation results. Infection of *A. thaliana* T-DNA mutant lines for eight of these factors revealed that genetic knock-out of the Double-stranded RNA-Binding protein 2 (DRB2) leads to increased TRV accumulation. In addition, over-expression of this protein caused a dramatic decrease in the accumulation of four unrelated plant RNA viruses, indicating that DRB2 has a potent and wide-ranging antiviral activity. We therefore propose B2:GFP-mediated pull down of dsRNA to be a novel and robust method to explore the proteome of VRCs *in planta*, allowing the discovery of key players in the viral life cycle.

**AUTHOR SUMMARY:** Viruses are an important class of pathogens that represent a major problem for human, animal and plant health. They hijack the molecular machinery of host cells to complete their replication cycle, a process frequently associated with the production of double-stranded RNA (dsRNA) that is regarded as a universal hallmark of infection by RNA viruses. Here we exploited the capacity of a GFP-tagged dsRNA-binding protein stably expressed in transgenic Arabidopsis to pull down dsRNA and associated proteins upon virus infection. In this manner we specifically captured short and long dsRNA from tobacco rattle virus (TRV) infected plants, and successfully isolated viral proteins such as the replicase, which attested to the successful isolation of virus replication complexes (VRCs). More excitingly, a number of host proteins, some of which have previously been involved in virus infection, were also captured. Remarkably, among a set of nine host candidates that were analyzed, eight showed dramatic re-localization to viral factories upon infection, and seven of these co-localized dsRNA-labeled VRCs. Genetic knock-out and over-expression experiments revealed that one of these proteins, *A. thaliana* DRB2, has a remarkable antiviral effect on four plant RNA viruses belonging to different families, providing ample validation of the potential of this experimental approach in the discovery of novel defense pathways and potential biotech tools to combat virus infections in the field. Being compatible with any plant virus as long as it infects Arabidopsis, we propose our dsRNA-centered strategy to be a novel and robust method to explore the proteome of VRCs *in planta*.

## INTRODUCTION

Viruses are obligate endocellular parasites that hijack their host’s molecular processes and machinery to multiply, a process that sometimes results in devastating diseases. Pivotal to a successful infection is the efficient replication of the viral genomic nucleic acid(s). In the majority of plant virus species the genome consists of one or more molecules of single-stranded positive polarity RNA, or (+)ssRNA. Replication is carried out by the virus-encoded RNA-dependent RNA-polymerase (RdRp), often part of a larger protein known as the replicase. This enzyme first copies the viral (+) genome into (-)ssRNA that is then used as a template for the production of progeny (+)ssRNA. Intrinsic to the RNA replication process is the generation of long double-stranded RNA (dsRNA) intermediates by the viral RdRp.

The replication of all known (+) strand RNA virus takes place on host membranes whose origin, whether the endoplasmic reticulum, chloroplasts, mitochondria, peroxisomes etc…, depends on virus species (reviewed in [1–4]). The progressive virus-induced recruitment of such membranes generally leads to dramatic reorganizations of the host endomembrane system into so called “viral factories”. These viral factories are the sites where all steps vital to the virus life are carried out including protein translation, RNA encapsidation and RNA replication *sensu stricto*. The specialized molecular entities on which RNA replication occurs within the viral factories are known as the virus replication complexes (VRCs). While a minimal VRC arguably consists of single- and double-stranded viral RNA and replicase, their precise composition, which depends on the virus and host species, remains largely unexplored. A number of studies have shown that specific host proteins can be integral part of these VRC and exert positive (pro-viral) or negative (anti-viral) effects on replication. Our knowledge on these host proteins is summarized in several exhaustive reviews [5–7]. These include among others RNA-binding proteins, RNA helicases, chaperones and proteins belonging to the RNA interference machinery (RNAi, or RNA silencing), which is the primary response to replicating viruses in plants and other eukaryotes [7, 8].

Antiviral RNAi against RNA viruses in the model plant *A. thaliana* is initiated by RNAse-III Dicer-Like enzymes DCL4 and DCL2, which cleave dsRNA into 21- and 22-nt small-interfering RNA (siRNA), respectively. These siRNA are then loaded into Argonaute (AGO) proteins, which use them as templates to recognize and cleave viral ssRNA in a sequence-specific manner. Viruses have evolved a vast array of strategies to evade or block RNAi, the best studied of which are viral suppressors of RNA silencing (VSRs). These proteins suppress silencing through a wide range of molecular strategies, from inhibition of dicing, to siRNA sequestration, to AGO degradation [9]. Of note, *A. thaliana* encodes several RNAse-III-like enzymes (RTLs) in addition to Dicers, but little is known regarding their function [10]. The accumulation of viral dsRNA *in planta* varies widely among (+)ssRNA virus and host species, as we have recently shown [11]. While the precise molecular events occurring during virus RNA replication are often unclear, it can be argued that rapid and efficient separation of the (+) and (-) RNA strands could not only allow more replication cycles to take place, but also constitute a powerful mechanism of RNA silencing evasion/suppression through removal of dsRNA.

A great deal of our knowledge on plant virus VRCs emerged from a series of seminal studies conducted with *Tomato bushy stunt virus* (TBSV) on yeast, a surrogate host used as a powerful biological tool to conduct genetic screens and functional studies on host factors involved in TBSV VRC activity (reviewed in [12]). While the authors of these studies, where possible, validated the results obtained in yeast and *in vitro* on plant species *N. benthamiana*, data obtained *in planta* on VRCs of other viruses remains sparse. Recent studies have reported methods to spatio-temporally visualize VRCs *in vivo* via fluorescently labeled dsRNA-binding proteins [8, 11, 13]. Candidate-based, reverse genetic approaches have been used to probe the involvement of host factors in VRC formation and activity in plants [14, 15]. These approaches, however, are necessarily based on prior discovery acquired by other experimental means. Another experimental strategy successfully used to characterize VRCs has been to pull down tagged viral proteins and analyze the resulting protein populations by mass spectrometry [16–18]. While this last method has provided valid and compelling data, we decided to investigate VRCs from a viral RNA-centered, rather than viral protein-centered, perspective. We hypothesized that pull-down of dsRNA from virus-infected plants followed by mass spectrometry of dsRNA-associated proteins would provide insight into the molecular composition of VRCs.

To test this hypothesis, we used the eGFP-tagged dsRNA-binding domain of FHV protein B2, which we have previously used to efficiently detect viral dsRNA *in vitro* and in constitutively-expressing *N. benthamiana* [11]. *Arabidopsis thaliana* being more appropriate for genetic studies than *N. benthamiana*, we first generated transgenic *A. thaliana* Col-0 plants stably expressing B2:GFP. We infected them with TRV, a well-studied (+)ssRNA virus, performed GFP immunoprecipitation and identified immunoprecipitated proteins by LC-MS/MS. In this manner, a number of virus- and host-encoded proteins could be identified. To validate their localization at B2:GFP-labeled TRV replication complexes, we tagged these candidates with tagRFP (tRFP hereafter) [19], transiently expressed them in healthy vs. TRV-infected 35S:*B2:GFP*/*N. benthamiana* [11] and observed the resulting leaf tissues by confocal laser scanning microscopy. We found that eight out of the nine *A. thaliana* proteins tested, including factors that have never before been linked to virus infection, re-localized strictly to B2:GFP-labeled TRV VRCs or alternatively to the larger TRV-induced viral factories. Among these candidates, DRB2, a dsRNA-binding protein was found to display antiviral activity. These results provide robust validation of dsRNA pull-down as an effective and high-throughput method for VRC characterization *in planta*. Furthermore, the results offer detailed snapshots of TRV replication complexes and viral factories, with host factors showing unique and distinct localization patterns in and around these complexes.

## RESULTS

### B2:GFP-mediated isolation of tobacco rattle virus dsRNA from *A. thaliana*

The double-stranded RNA-binding B2:GFP protein, when ectopically expressed in transgenic 35S:*B2:GFP*/*N. benthamiana*, has been previously shown to specifically associate with the VRCs of several positive-strand RNA viruses from plants and insects [11]. Following these findings, we wished to further exploit B2:GFP as a biochemical bait to explore the composition and biology of RNA VRCs, the pivotal element of which is dsRNA. To do so, and given the versatility of *A. thaliana* as a model plant species, we first produced homozygous 35S:*B2:GFP* transgenic plants. Although in this work we focused essentially on the 35S:*B2:GFP/*Col-0 line, 35S:*B2:GFP* was also introduced into various genetic backgrounds including mutants of the core antiviral Dicer-Like genes, *dcl2-1*, *dcl4-2* and triple *dcl2-1/dcl3-1/dcl4-2* (**Supplementary Figure 1A,B,C**). The rationale behind this choice is that DCL proteins are arguably the best-known RNAse III enzymes in plants, and the small-interfering RNA (siRNA) they generate from virus-derived dsRNA precursors are the effectors of RNA silencing, the main antiviral defense in plants [9, 20]. Similarly to the B2:FP *N. benthamiana* lines [11] and despite the clear expression of B2:GFP (**Supplementary Figure 1A),** the different lines showed little to moderate developmental phenotypes (**Supplementary Figure 1C**). These were very reminiscent of - but distinct from - those caused by ectopic expression of other RNA silencing suppressors such as P19 or HC-Pro [21, 22]. Such phenotypes may be determined (i) by inhibition of long dsRNA processing into siRNA or (ii) by disruption of miRNA function through their sequestration, or both.

The full-length FHV B2 protein has been shown to be a suppressor of RNA silencing [23, 24], and proposed to act through both inhibition of dicing and sequestration of siRNA [25]. To investigate whether the GFP-tagged 73 amino acid dsRNA binding domain of B2 that lacks the residues involved in the interaction with PAZ domains of Dicer proteins [26, 27] also acts as a suppressor of RNA silencing, we performed a standard GFP silencing patch test on *N. benthamiana* leaves (**Supplementary Figure 1D,E**). B2, as a C-terminal fusion to tRFP (B2:tRFP) was able to suppress silencing of the GFP transgene, as was turnip crinkle virus suppressor P38. By contrast, a C44S, K47A double-mutated version of B2:tRFP impaired in dsRNA binding [11] was unable to suppress silencing, suggesting that suppression activity is dsRNA-binding-dependent and likely DCL-binding-independent. Next, we investigated the effects of stably expressed B2:GFP on endogenous small RNA pathways: biogenesis of microRNAs 159 and 160 was not perturbed in 35S:*B2:GFP*/Col-0 plants, while biogenesis of siRNA such as endo-siRNA (IR71) and *trans*-acting siRNA (TAS1) was completely abolished (**Supplementary Figure 1B**). Whether these defects in endo-siRNA and *trans*-acting siRNA biogenesis are responsible of the observed developmental phenotypes remains to be determined.

We then infected 35S:*B2:GFP*/Col-0 plants with a recombinant TRV carrying part of the phytoene desaturase (PDS) gene [28]. As expected, the control Col-0 plants showed minor viral symptoms and the typical bleaching phenotype linked to PDS gene silencing. In contrast, the B2:GFP-expressing plants showed no significant leaf discoloration but severe viral symptoms (**Figure 1A**) and death of the plants occurred before flowering (not shown), well in agreement with the efficient RNA silencing suppression activity of the B2 dsRNA-binding domain. Observation of systemically infected leaves by confocal microscopy showed TRV-induced re-localization of B2:GFP to distinct cytosolic mesh-like structures (**Figure 1B**) very reminiscent to those observed in 35S:*B2:GFP*/*N. benthamiana* and shown to correspond to TRV-induced VRCs [11]. Northern analysis of RNA from TRV-PDS systemically infected 35S:*B2:GFP*/*N. benthamiana* and 35S:*B2:GFP*/Col-0 revealed that B2:GFP caused a striking over-accumulation of viral (+)ssRNA in both plant species (**Figure 1C**). This is also in agreement with B2:GFP activity as a suppressor of RNA silencing, and could be recapitulated in a *dcl2-1/dcl4-2* double mutant (*dcl24*), which lacks the two main antiviral Dicers (**Supplementary Figure 1F**)[29]. B2:GFP also caused a tremendous increase in long double-stranded RNA content, likely corresponding to replication intermediates, as determined by northwestern blotting in both B2:GFP-expressing Col-0 and *N. benthamiana* plants (**Figure 1D**). In contrast, TRV-derived siRNA were differentially accumulated between Arabidopsis and *N. benthamiana*. Thus, while B2:GFP expression led to an overall reduction in 21 and 22 nt vsiRNA species in Col-0 plants, the opposite effect was observed in *N. benthamiana* (**Figure 1E**). Conversely, miR159 and U6-derived snRNA accumulation was unaffected by the presence of B2:GFP in both plant species (**Figure 1E**, **Supplementary Figure 1B**). These results suggest that B2:GFP interferes strongly with TRV RNA processing by DCL enzymes, thereby promoting viral replication.

**Figure 1:**
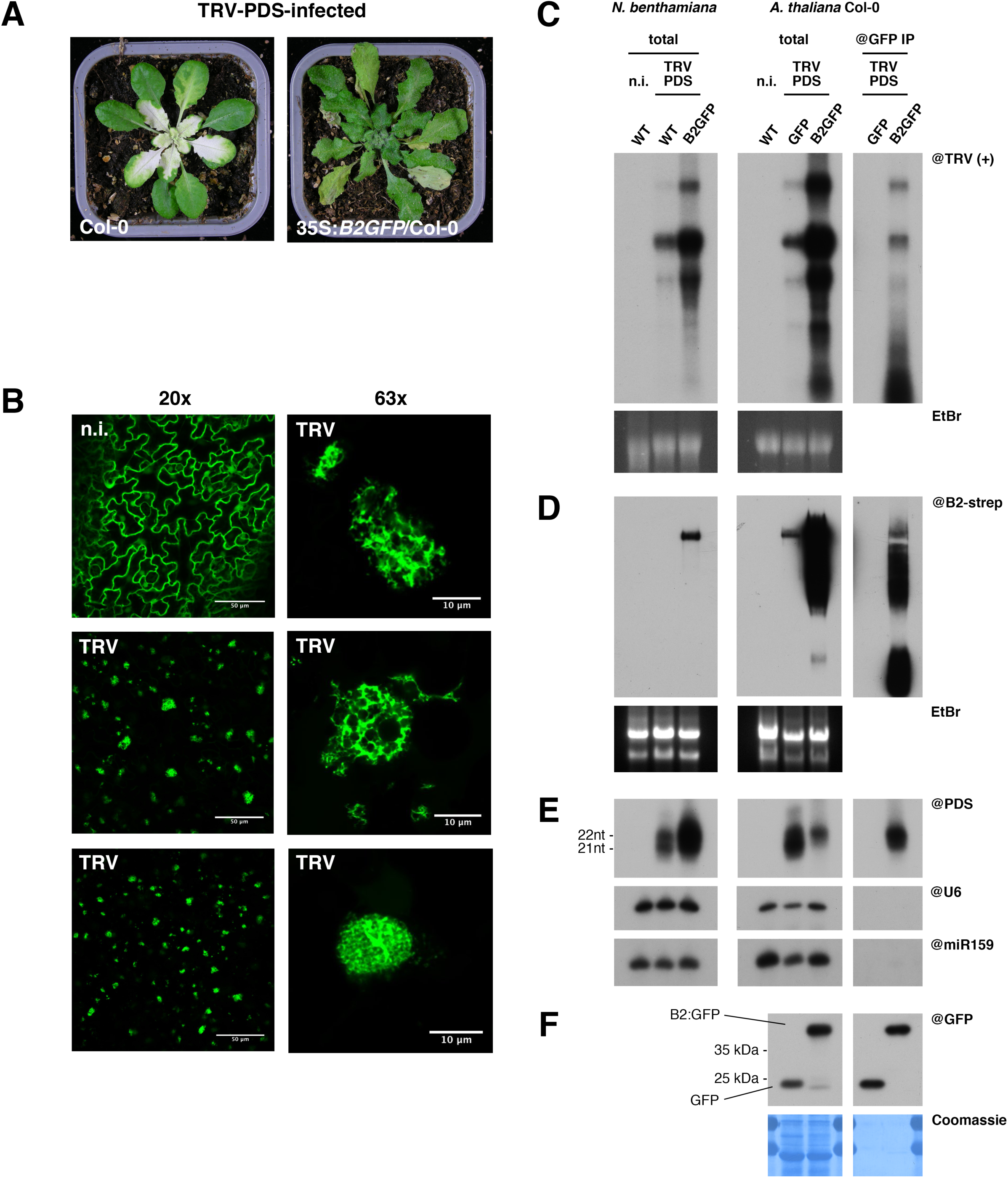
Immunoprecipitation of B2:GFP allows the isolation of TRV dsRNA *in vivo*. **(A)** Photos of *A. thaliana* Col-0 and 35S:*B2:GFP*/Col-0 plants 13 days post-infection with TRV-PDS. **(B)** Confocal microscopy analysis of non-infected (n.i. - top left) and TRV-PDS systemically-infected leaves of 35S:*B2:GFP*/Col-0 plants. On the right, higher magnification (63x) images of TRV replication complexes from the same tissues as those visible at lower magnification (20x) on the left middle and bottom. **(C)** Northern blot analysis of high molecular weight RNA from total fractions of TRV-PDS-infected wild-type or 35S:*B2:GFP N. benthamiana* (left), and from total (middle) and anti-GFP immunoprecipitated (right) fractions from infected 35S:*GFP* and 35S:*B2:GFP/*Col-0 *A. thaliana*. **(D)** Northwestern blot analysis of native-state high molecular weight RNA from samples described in (C). EtBr staining was used as loading control in (C) and (D). **(E)** Northern blot analysis of low molecular weight RNA from samples described in (C). The probes were applied sequentially on the same membrane in successive rounds of probing and stripping. snU6 and miR159 were used as loading controls. **(F)** Western blot analysis of proteins from the same immunoprecipitation experiment analyzed in (C). Coomassie staining was used as loading control. Source data is available with the Blotting Source Data.

As a first experiment establishing B2:GFP as a tool to study VRC composition, we performed anti-GFP immunoprecipitations (IPs) from TRV-PDS-infected 35S:*B2:GFP*/Col-0 plants and analyzed their composition in (+)strand viral RNA (**Figure 1C**), long dsRNA (**Figure 1D**), siRNA (**Figure 1E**) and proteins (**Figure 1F**). As a negative control, we included TRV-PDS-infected 35S:*GFP*/Col-0 plants. Northern and northwestern analyses performed on IPed RNA revealed that immune complexes contained (+)ssRNA, long dsRNA and 22nt vsiRNA, but no U6 and miR159 (**Figure 1C-E**). Interestingly, and in contrast with our previous report *in vitro* [11] but well in agreement with the capacity of B2 to bind dsRNAs longer than 18 bp [25], antiviral siRNA were immunoprecipitated (**Figure 1E**). Western analysis of proteins from the same experiment revealed efficient IP of both GFP and B2:GFP (**Figure 1F**). Altogether we concluded that immunoprecipitation allowed the isolation of TRV double-stranded replication intermediates, the core element of VRCs.

### Immunoprecipitation and identification of B2:GFP-associated viral and host proteins by mass spectrometry

Once established that immunoprecipitation of B2:GFP from plants allowed efficient isolation of virus replication dsRNA intermediates, we wondered whether these complexes contain specific virus- and host-encoded proteins. To address this question we performed anti-GFP IP on TRV-PDS-infected 35S:*GFP*/Col-0 vs. 35S:*B2:GFP*/Col-0 in triplicate, and analyzed the immunoprecipitated proteins by mass spectrometry (MS). The complete list of identified viral and host proteins from this analysis in shown in **Supplementary Table 1**. We also performed the same IP and MS analysis on non-infected 35S:*GFP*/Col-0 vs. 35S:*B2:GFP*/Col-0 plants (**Supplementary Table 2**).

A preliminary analysis by immunoblot confirmed efficient and reproducible B2:GFP and GFP immunoprecipitation (**Supplementary Figure 2**), which could be confirmed by MS, reads from B2:GFP and GFP being the most abundant (**Figure 2A,B, Supplementary Table 1**). We next searched and ranked accessions that were identified only - or highly enriched - in 35S:*B2:GFP*/Col-0 samples (**Figure 2A,B**, **Supplementary Table 1**). As expected, the TRV replicase (Uniprot accession Q9J942) was among the most abundant proteins detected in immunoprecipitates from B2:GFP samples (**Figure 2**). This result, along with the previously described detection of viral dsRNA in analogous IPs (**Figure 1**), suggests that B2:GFP immunoprecipitation allows the isolation of TRV VRCs. This hypothesis is further supported by the detection of TRV coat protein (CP, Uniprot Q88897) and 16k suppressor of silencing (Uniprot Q77JX3) in B2:GFP immunoprecipitates (**Figure 2**). It also suggests that the 16k and CP associate directly or indirectly to dsRNA. Although unlikely, we can’t at this point rule out that one or more of these TRV proteins bind B2:GFP and not dsRNA. In addition to TRV-encoded proteins, MS analysis allowed also the identification of 110 host proteins exclusively present in immunoprecipitates from B2:GFP-expressing plants (**Figure 2** and **Supplementary Table 1**), which we considered as replication complex-associated host protein candidates. 29 of these proteins were significantly enriched in the IPs with an adjusted p-value < 0.05 (**Figure 2B**).

**Figure 2:**
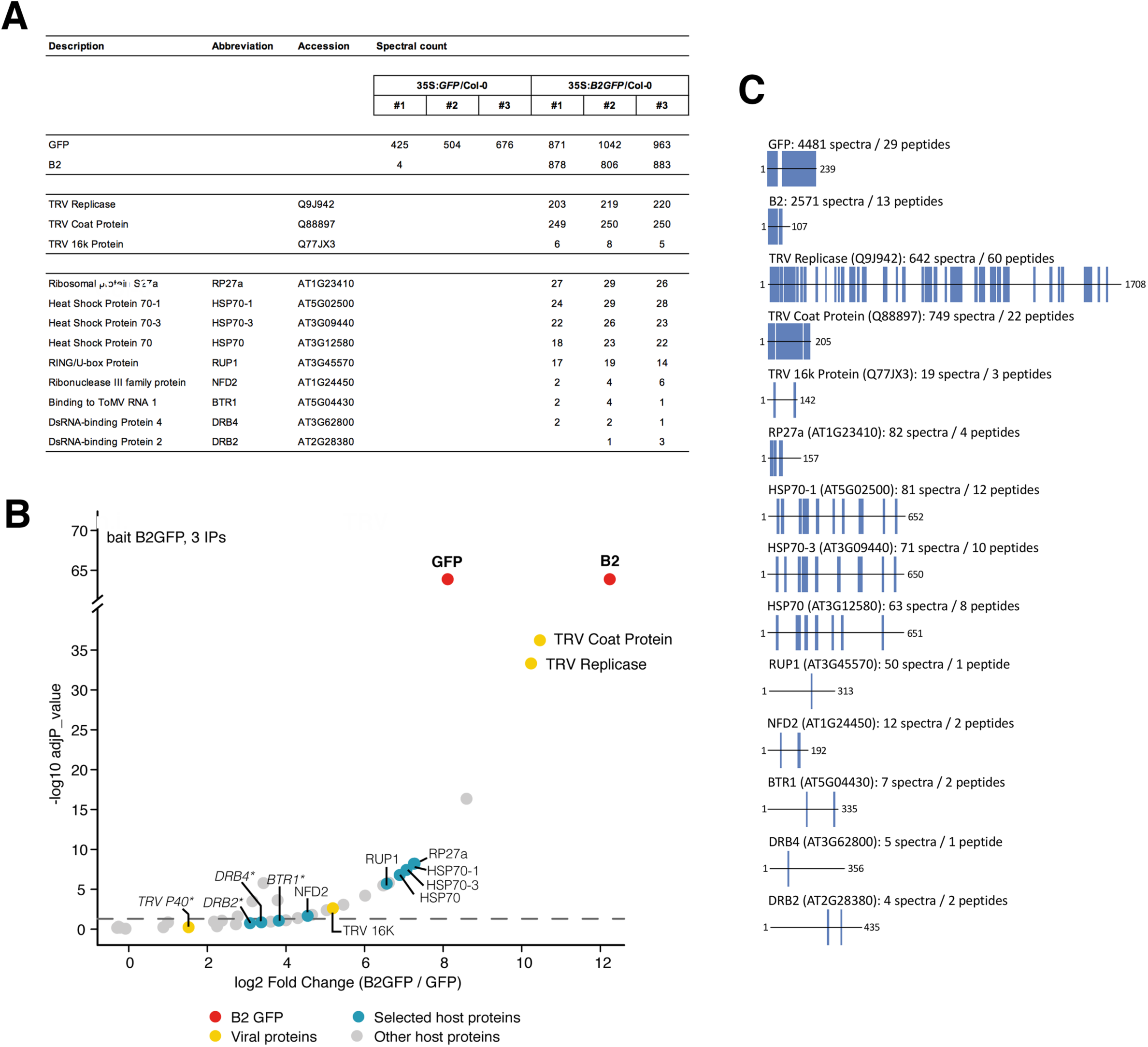
Immunoprecipitation of B2:GFP allows the isolation of TRV proteins and host factors. Mass spectrometry analyses of anti-GFP immunoprecipitated proteins from TRV-PDS-infected 35S:*GFP*/Col-0 and 35S:*B2:GFP*/Col-0 plants. Three technical replicates were performed and analyzed per genotype. *A. thaliana* proteins shown here are the ones that have been tested in this study. Abbreviations correspond to those available in literature, except for RP27a and RUP1, which have been here assigned for lack of previously established ones. Accession numbers correspond to UniProt (TRV proteins) and TAIR (*A. thaliana* proteins) databases. **(A)** Table containing the spectral counts obtained per technical replicate for bait proteins (GFP + B2), TRV proteins and *A. thaliana* proteins. **(B)** Volcano plot representation shows the enrichment of proteins from TRV-infected plants that co-purified with B2:GFP. Y- and X-axis display adjusted p-values and fold changes, respectively. The dashed line indicates the threshold above which proteins are significantly enriched (adjP □ < □ 0.05). The source data are available in **Supplementary Table 2**, (C) Sequence coverage obtained on all the proteins listed in (A). For each of the 14 entries, the length of the protein is displayed while the covered residues are highlighted with blue vertical bars. The total number of spectra matching on each protein in the three replicates is indicated, as well as the corresponding total number of unique peptide sequences. The complete protein list can be found in Supplementary Table 2.

With the aim to evaluate the association of candidate host proteins to replication complexes and considering their high number (**Supplementary Table 1**), we first established a priority list of nine *A. thaliana* gene products that were either detected with high spectral counts (AT5G02500, AT3G09440, AT3G12580) or confirmed/potential RNA-binding/interacting proteins from literature or NCBI annotation (AT1G24450, AT5G04430 [30], AT3G62800 and AT2G28380 [31]), or both (AT1G23410, AT3G45570)(**Figure 2A,B**). The distribution of peptide reads along these selected proteins, along with the TRV and bait proteins, is shown in **Figure 2C**. It should be noted that four of the candidates (AT1G23410, AT3G09440, AT5G02500 and AT3G12580) were also present in B2:GFP IPs from non-infected plants (**Supplementary Table 2**) which may reflect their dsRNA binding activity in both healthy and virus-infected plants. However, the spectral count of peptides from these proteins was a fraction of that detected in IPs from TRV-infected plants, despite the spectral counts of the bait proteins being comparable. All other candidates were not detected in B2 IPs from non-infected plants. Finally, we excluded from our priority list a number of proteins that were significantly enriched in B2:GFP vs. GFP plants due to the number of candidates to analyze and their apparent lack of significance in viral replication process based on literature. This includes for instance the most enriched protein, a myrosinase present in Brassica crops with anti-microbial activity and involved in defense against herbivores [32].

### tRFP does not label TRV replication complexes *in planta*

In a second step, we tested the subcellular localization of the selected candidates in relation to B2:GFP in healthy and TRV-infected plants. To do so, we opted for the 35S-driven transient expression of the Arabidopsis candidates as N- or C-terminal fusions to tRFP in healthy or TRV-infected 35S:*B2:GFP*/*N. benthamiana*. In all cases, confocal imaging was performed 3-4 days post agro-infiltration, a time that was found to be optimal for TRV-infection and transient expression of protein candidates.

As an absolute prerequisite to our validation pipeline of candidate proteins and considering tRFP was used as reporter tag, we first carefully analyzed the intracellular distribution of tRFP with respect to TRV replication complexes in 35S:*B2:GFP*/*N. benthamiana*. As expected, tRFP as well as B2:GFP showed a typical nucleo-cytoplasmic localization in cells from healthy plants (**Figure 3A**), well in agreement with our previous report using the same experimental system [11]. Crucially, upon infection the intracellular distribution of tRFP remained unchanged, while B2:GFP concentrated to bright cytoplasmic cotton-ball-like structures often adjacent to the nucleus (**Figure 3B**). These large structures were previously shown to correspond to TRV viral factories enriched in mitochondria-derived membranes [11] on which replication of TRV is thought to occur [33]. Importantly, our data clearly show that while B2:GFP is highly enriched in TRV replication factories, tRFP alone is significantly depleted from these structures (**Figure 3B**), in agreement with the behavior of tRFP as cytoplasmic and validating tRFP as a reporter protein with which to tag the candidates of interest.

**Figure 3:**
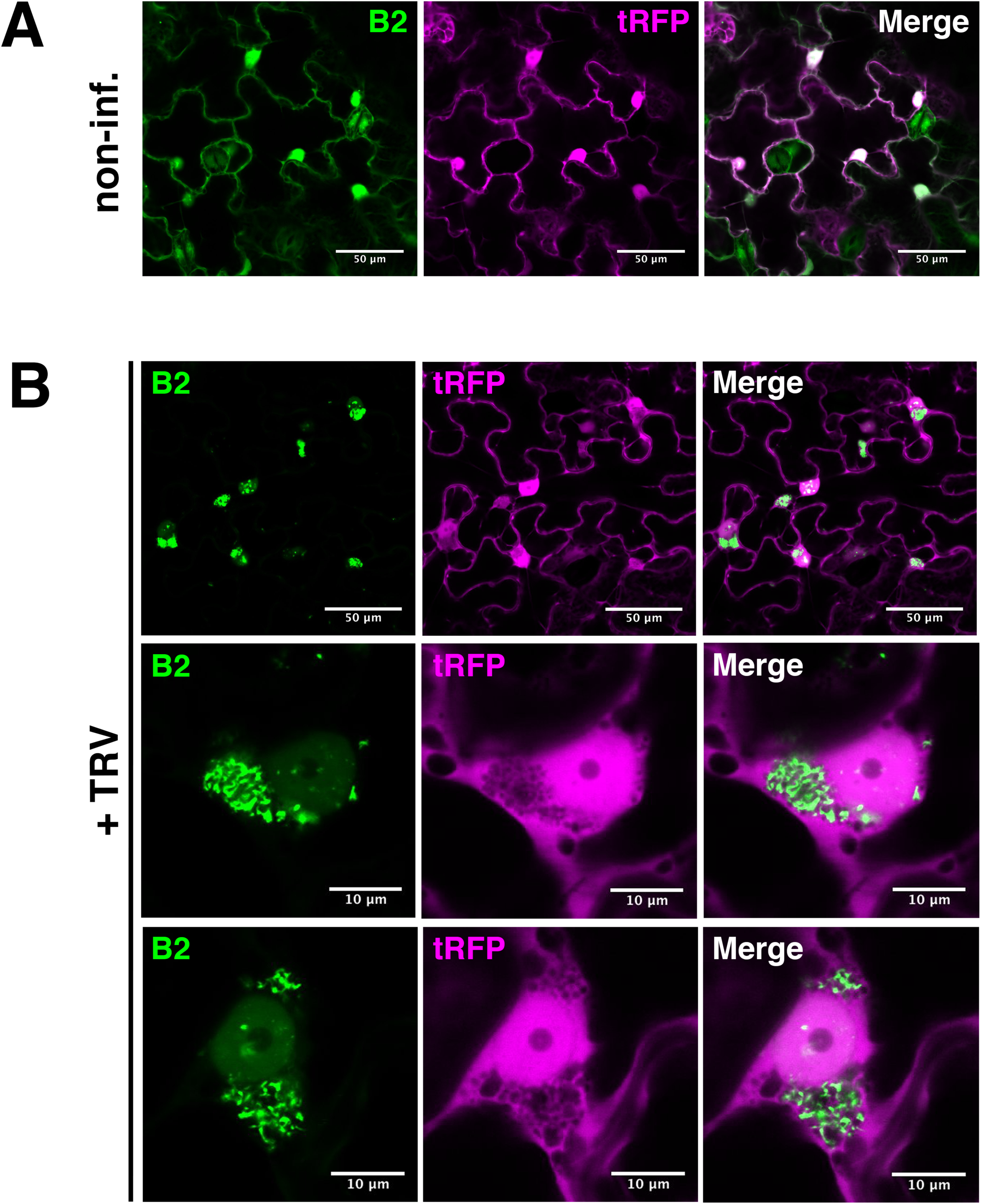
tRFP does not re-localize to TRV replication complexes. Laser confocal microscopy on 35S:*B2:GFP/N. benthamiana* leaves transiently expressing 35S:*tRFP*. **(A)** Acquisition from non-infected leaf disks (20x objective). Scale bars indicate 50 μm. **(B)** Acquisition of TRV-PDS-infected leaf disks (20x objective, top), focused on TRV replication complexes (63x objective, middle and bottom). Scale bars indicate 50 and 10 μm, respectively. Additional acquisitions can be found with the Microscopy Source Data.

### Double-stranded RNA-binding proteins (DRBs) perfectly colocalize with B2-labeled viral replication complexes

DRBs are proteins with dual dsRNA-binding motifs with five representatives in the Arabidopsis genome [31, 34]. Despite showing low spectral counts in our IPs, two DRBs were identified in our analysis: DRB2 (AT2G28380, total counts: 4, **Figure 2A**) and DRB4 (AT3G62800, total counts: 5, **Figure 2A**) that were obvious candidates to test (**Figure 4**).

**Figure 4:**
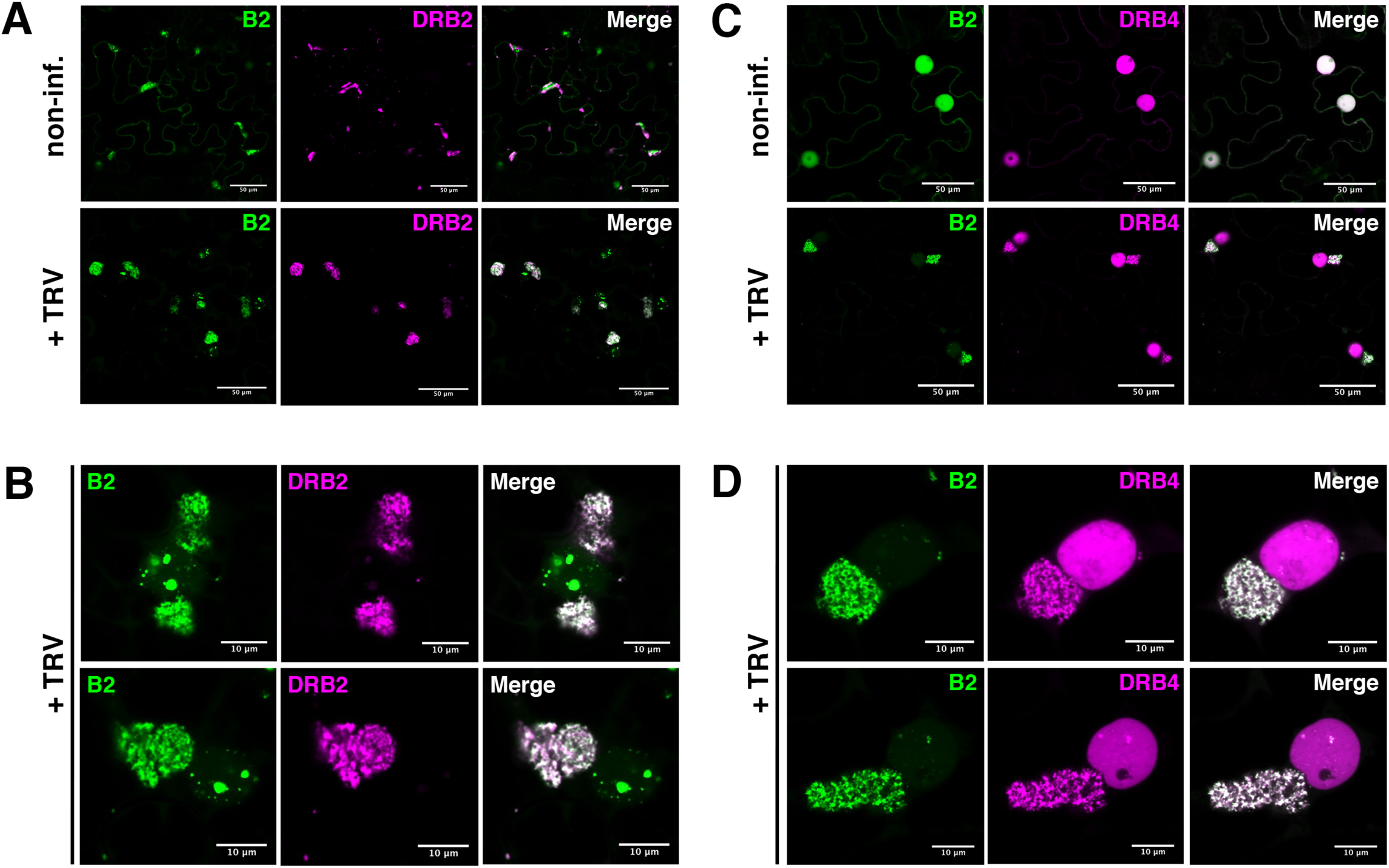
*A. thaliana* double-stranded RNA-binding proteins localize at TRV replication complexes. Laser confocal microscopy on 35S:*B2:GFP/N. benthamiana* leaves transiently expressing 35S:*DRB2:tRFP* (A,B) or 35S:*DRB4:tRFP* (C,D). **(A)** Acquisitions with 20x objective of non-infected (top) and TRV-PDS-infected (bottom) leaf disks expressing DRB2:tRFP. Scale bars indicate 50 μm. **(B)** Acquisitions with 63x objective of TRV-PDS-infected leaf disks of tissue described in (A). Scale bars indicate 10 μm. **(C,D)** As in (A,B), but from tissue expressing DRB4:tRFP. Additional acquisitions can be found with the Microscopy Source Data.

DRB2 was recently shown to localize to the replication complexes of different RNA viruses [8], to be able to bind dsRNA [35] and to play a role in endogenous small RNA biogenesis [31, 36, 37]. In non-infected plants DRB2:tRFP and B2:GFP localized to partially overlapping cytoplasmic and nuclear structures. Interestingly, over-expression of DRB2 changed the localization pattern of B2 from a predominantly nuclear localization (**Figure 3A** and [11]) to DRB2-labeled cytoplasmic structures as if B2:GFP was recruited to DRB2 localization sites (**Figure 4A**). Remarkably, such redistribution of B2 was not observed upon overexpression of DRB4 (**Figure 4C**). Crucially, near-perfect colocalization of DRB2:tRFP and B2:GFP was observed in the VRCs upon TRV-PDS infection (**Figure 4A**), which was particularly evident at high magnification (**Figure 4B**). Moreover, while DRB2:tRFP was almost exclusively found in the cytoplasmic VRCs upon infection, a substantial fraction of B2:GFP remained associated to nuclear structures likely containing dsRNA (**Figure 4B**). This suggests that although both proteins are susceptible to bind dsRNA, their intracellular targeting is likely not exclusively dsRNA-dependent.

DRB4 has been shown to be both a co-factor of DCL4 in small RNA biogenesis and an inhibitor of DCL3 in endogenous inverted-repeat RNA processing in *A. thaliana* [38, 39]. More relevantly here, DRB4 is involved in the defense against RNA viruses [40, 41]. When we expressed DRB4:tRFP in non-infected tissue, this protein accumulated predominantly to the nucleus where it colocalized with B2:GFP (**Figure 4C**), in agreement with previous reports [8, 11]. Upon TRV infection DRB4:tRFP was clearly redistributed to VRCs, where it perfectly colocalized with B2:GFP (**Figure 4C,D**). In contrast to DRB2 that was barely detectable in the nucleus (**Figure 4B**), a significant fraction of DRB4:tRFP remained nuclear upon infection (**Figure 4C,D**).

Altogether, the robust colocalization of both double-stranded RNA-binding proteins DRB2 and DRB4 with B2:GFP during infection provide (i) further evidence that the TRV-viral factories are indeed cytoplasmic dsRNA hotspots and, more importantly, (ii) a first validation of the immunoprecipitation procedure.

### Proteins previously linked to viral infection localize at/near VRCs

A family of proteins that emerged with high spectral counts were those belonging to the HSP70 family: HSP70 (AT3G12580, 63 counts), HSP70-1 (or HSC70-1, AT5G02500, 81 counts) and HSP70-3 (or HSC70-3, AT3G09440, 71 counts) (**Figure 2A**). Members of this family of chaperones have been shown in several studies to play key roles in virus infection cycles (reviewed in [5, 42]). They can regulate viral life cycles both positively and negatively, and depending on the virus, they affect VRC formation, virus movement and coat protein homeostasis, among other processes. Three recent studies showed that unrelated plant viruses hijack HSP70 to greatly enhance virus replication [43–45].

All three HSP70 members were tested in TRV-infected and non-infected 35S:*B2:GFP*/*N. benthamiana* **(Figure 5 and Supplementary Figure 3)**. When overexpressed in healthy plants HSP70:tRFP, HSP70-1:tRFP, HSP70-3:tRFP localized essentially to distinct cytoplasmic foci whose number, size and distribution were specific for each of the three HSP70 observed (**Supplementary Figure 3**). Remarkably, upon infection, HSP70-1:tRFP **(Figure 5B)** and HSP70-3:tRFP **(Figure 5C)** were clearly redistributed to TRV viral factories enriched in B2:GFP. In contrast, the localization pattern of HSP70:tRFP remained essentially unaffected upon infection, with no obvious colocalization of B2:GFP with HSP70:tRFP-labeled foci **(Figure 5A, Supplementary Figure 3A)**.

**Figure 5:**
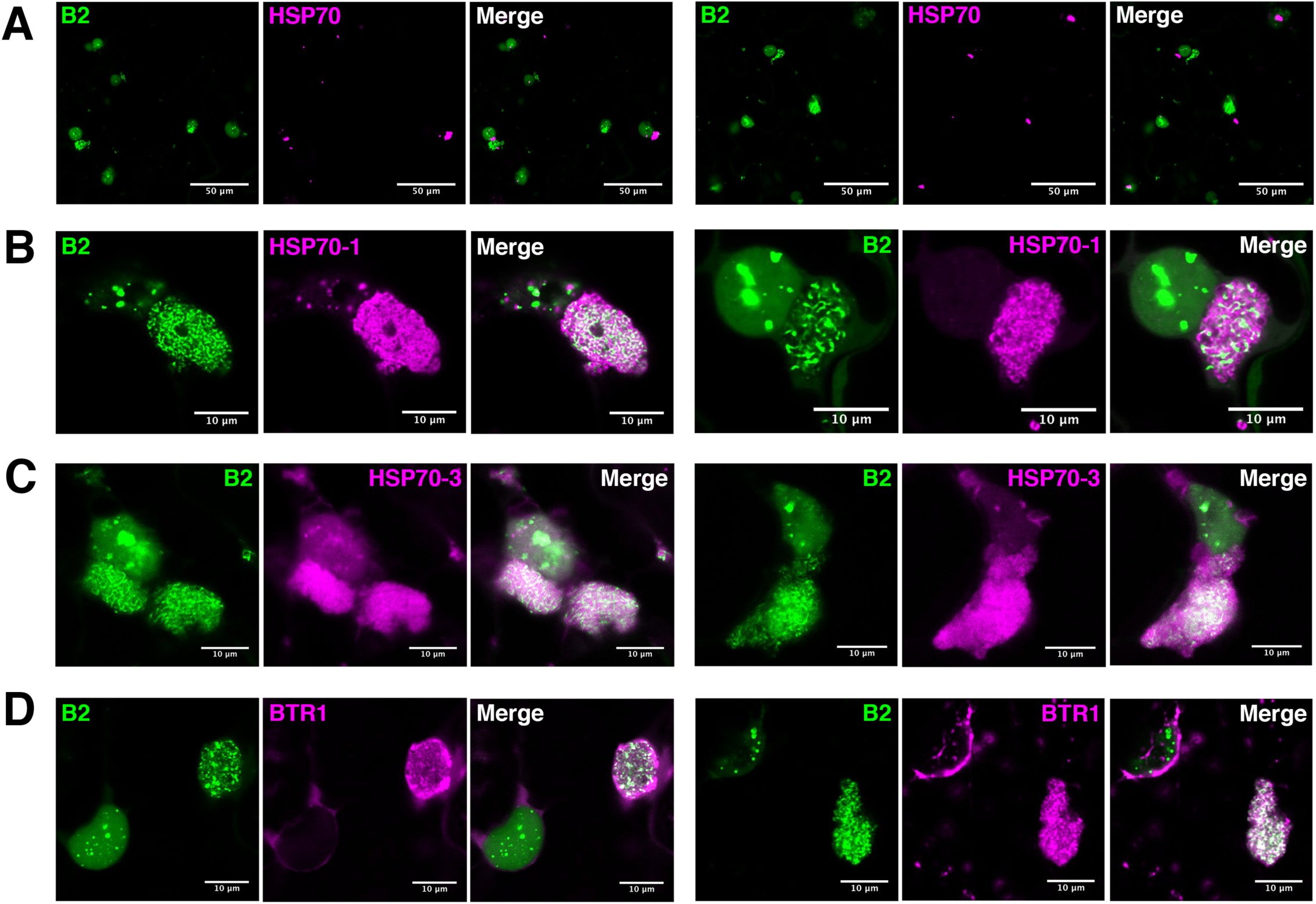
Proteins previously implicated in viral life cycle localize at or near the replication complexes. Laser confocal microscopy on 35S:*B2:GFP/N. benthamiana* TRV-PDS-infected leaf disks transiently expressing **(A)** 35S:*HSP70:tRFP,* **(B)** 35S:*HSP70-1:tRFP*, **(C)** 35S:*HSP70-3:tRFP,* **(D)** 35S:*BTR1*:*tRFP*. Acquisitions in (A): 20x objective, scale bars indicate 50 μm. Acquisitions in (B, C, D): 63x objective, scale bars indicate 10 μm. Additional acquisitions can be found with the Microscopy Source Data.

It should be noted that despite the clear redistribution of HSP70-1:tRFP and HSP70-3:tRFP upon infection, only partial colocalization was detected between these proteins and B2:GFP **(Figure 5B,C)**. The latter appeared engulfed in large HSP70-1 or HSP70-3-containing bodies, likely corresponding to larger viral factories. This sub-localization is in sharp contrast with the near perfect colocalization of B2:GFP with DRB2 and DRB4 upon infection (**Figure 4**). Altogether our results suggest that HSP70-1 and HSP70-3, contrarily to HSP70, are components of the TRV viral factories. However, contrarily to B2, DRB2 and DRB4 that directly interact with dsRNA, HSP70-1 and HSP70-3 are likely involved in indirect interactions with TRV replication complexes, perhaps via the TRV replicase or other viral or host components. Interestingly, it has recently been observed using the B2:GFP system that dsRNA-containing VRCs constitute only a part of the structures induced by PVX, which in fact also contain viral ssRNA and coat protein [11]. The components and activities harbored within these larger “viral factories” are still largely unknown, but the localization patterns of HSP70-1 and HSP70-3 suggests that these proteins associate not only to replication complexes but also to other entities within viral factories.

Next, we tested the localization of an RNA-binding protein present in our IP MS list that was previously shown to associate to plant virus RNA. This protein, known as Binding to ToMV RNA (BTR1, AT5G04430, 7 counts, **Figure 2A**), was identified through affinity purification of tagged viral RNA and found *in vitro* to bind to the 5’ region of the (+) polarity RNA of ToMV, a tobamovirus [30]. In our experimental system, BTR1:tRFP localized to numerous cytoplasmic punctate structures at the cell periphery in non-infected cells **(Supplementary figure 3D)**. Upon TRV-PDS infection, BTR1 was seen to clearly localize at B2:GFP-labeled VRCs, while to some extent maintaining the localization visible in non-infected cells **(Supplementary figure 3D)**. At high magnification it is possible to see that BTR1 did not strictly and exclusively colocalize with B2:GFP, but could also be seen in the areas surrounding B2:GFP-labeled dsRNA hotspots (**Figure 5D**). Similarly to HSP70-1 and HSP70-3, it is possible that BTR1 associates not only to VRCs but also to other entities within viral factories.

### Novel proteins are localized at/near replication complexes

Among the potential TRV replication complex-associated proteins identified through IP, we tested three on which we found no specific function in virus process from the literature: a RING/U-box protein (AT3G45570, 50 counts) and Ribosomal Protein S27a (AT1G23410, 82 counts) and NFD2 (Nuclear Fusion Defective 2 – AT1G24450, 12 counts, **Figure 2A**).

The RING/U-box protein, which we will refer to as RUP1, belongs to the E3 ubiquitin ligase RBR family. The N-terminal half of the protein is homologous to the RNAse H superfamily, followed on the C-terminal half by a RING-type zinc-finger domain and an IBR (In Between Ring fingers) domain. The Ribosomal Protein 27a, here abbreviated to RP27a, is a small protein of 156 aminoacids with a ubiquitin domain N-terminal half and a zinc-binding ribosomal protein superfamily C-terminal section. Importantly, all RP27a peptides detected in B2:GFP co-immunoprecipitated samples belong to the ubiquitin domain, and also match with the protein sequences of UBQ1 through UBQ14 (**Supplementary Figure 5**). This suggests ubiquitination of one or more proteins present in the immunoprecipitates that possibly associated to TRV replication complexes. To investigate which protein/s were ubiquitinated, we searched the mass spectrometry dataset for di-glycine footprints, a hallmark of ubiquitination. Interestingly, the only protein found to contain such a feature was RP27a itself, only on lysine-48 (**Supplementary Figure 5**), suggesting self-ubiquitination and/or the formation of lysine-48 polyubiquitin chains. Given that no other di-glycine footprint was found in our spectrometry dataset, the proteins targeted by these chains may have been below detection level and remain to be identified. Finally, NFD2 was first identified as a factor involved in karyogamy, the fusion of polar nuclei within the central cell of the female gametophyte prior to fecundation and the fusion of the sperm cells’ nuclei with the egg cell and the central cell upon fecundation [46]. This protein, containing an RNAse III domain, has been also described as RNAse Three-Like 4 (RTL4) [10].

In healthy cells, tRFP-tagged RUP1 and RP27a displayed a nucleo-cytoplamic distribution (**Supplementary Figure 4A,B)**, while NFD2 was essentially found in numerous cytoplasmic bodies (**Supplementary Figure 4C)**. Upon TRV infection, all three proteins were clearly re-localized to or near B2:GFP-labeled dsRNA hotspots (**Figure 6**, **Supplementary Figure 4).** More precisely, RUP1 and RP27a showed patterns similar to those observed with HSP70-1, with extensive overlap with B2:GFP as seen from the white color in the merged panels (**Figure 6A,B**). This suggests that RUP1 and RP27a associate not only with VRCs but also to other entities within viral factories. Interestingly, NFD2 showed a pattern of localization different from the other proteins tested in this work (**Figure 6C, Supplementary Figure 4C)**. Although localization of NFD2 and B2 seemed mutually exclusive, NFD2 being absent from B2:GFP-labeled structures and vice versa, a continuum between B2:GFP-labeled hotspots and NFD2-labeled structures was observed (**Figure 6C**), suggesting that NFD2 is intimately linked to TRV-induced subcellular entities and was therefore immunoprecipitated. In addition, the complete localization of NFD2 in close proximity to TRV replication complexes was in stark contrast with the perinuclear and cytoplasmic point-form localization of NFD2 in non-infected plants **(Supplementary Figure 4C**).

**Figure 6:**
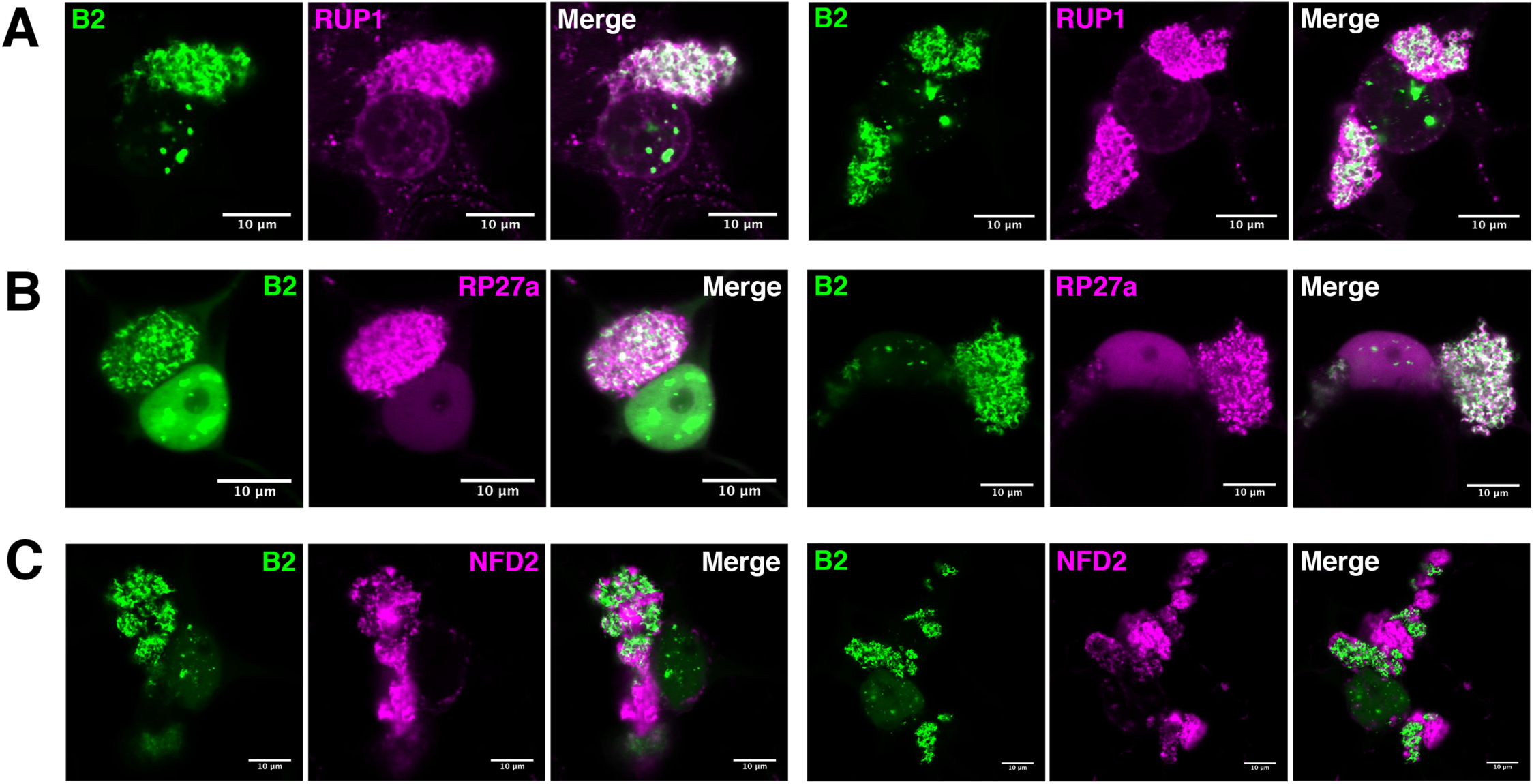
Localization of previously undescribed proteins at or near the replication complexes. Laser confocal microscopy (63x objective) on 35S:*B2:GFP/N. benthamiana* TRV-PDS-infected leaf disks transiently expressing **(A)** 35S:*RUP1*:*tRFP*, **(B)** 35S:*tRFP:RP27a*, **(C)** 35S:*tRFP:NFD2*. Scale bars indicate 10 μm. Additional acquisitions can be found with the Microscopy Source Data.

### Knock-out of DRB2 potentiates TRV systemic infection in a Dicer-independent manner

Next, we tested whether genetic knock-out of the candidate genes analyzed would lead to changes in TRV systemic accumulation. To do so, we acquired *A. thaliana* lines with T-DNA insertions in the genes of interest: *drb2-1*, *drb4-1* and *drb2-1/drb4-1* [47], *btr1-1* [30], SALK_078851 (*rup1*), SALK_093933 (*rp27a*), SALK_088253 (*hsp70*), SALK_135531 (*hsp70-1*) and SALK_020290 (*hsp70-3*). No lines carrying insertions in the annotated 5’UTR or coding sequence of *NFD2* were found. We infected ten plants per genotype and harvested the inoculated leaves 3 days post-infection (dpi), dividing the plants of each genotype into two equal pools. In parallel, an identical set of infections was performed, and systemically infected leaves were harvested 12 dpi. Northern blot analysis of the total RNA from these samples revealed that, while there were no noticeable changes in TRV RNA accumulation in the inoculated leaves at 3dpi (**Figure 7A**), both the single *drb2-1* and double *drb2-1/drb4-1* mutants showed markedly increased TRV accumulation in systemic leaves compared to control Col-0 plants at 12 dpi (**Figure 7B**). This suggests that DRB2 could play an antiviral function with respect to TRV. A moderate increase in TRV accumulation was also observed in systemic leaves of *btr1-1* mutants, in agreement with published work on ToMV [30] (**Figure 7B**). Considering that major differences in TRV accumulation between SALK lines and Col-0 control were essentially restricted to *drb2-1* and *drb2-1/drb4-1* lines, we decided to focus our attention on possible antiviral function of DRB2. Since DRB proteins have been shown by several studies to be involved in small RNA biogenesis [35, 37, 38, 39, 40] and DRB2 genetic knock-out has been shown to impact accumulation of several microRNAs [36], we decided to analyze the viral siRNA (vsiRNA) present in the *drb* mutants described above (**Figure 7C**). Northern blot analysis of small RNAs revealed an increase in vsiRNA in the *drb2-1* and *drb2-1/drb4-1* mutants analyzed. This most likely reflects the increase in TRV genomic RNAs in these samples (**Figure 7B**), which are substrates for vsiRNA biogenesis. Moreover, DRB2 knock-out didn’t cause any noticeable changes in the vsiRNA size distributions. These observations, overall, lead us to conclude that knock-out of DRB2 (i) positively impacts TRV RNA and vsiRNA steady-state levels and (ii) does not cause changes in the respective contributions of DCL2, DCL3 and DCL4 to this process. These observations are in line with what has been previously observed for TuMV and TSWV [47]. Therefore, the increase of TRV systemic accumulation observed in *drb2* mutants is likely not due to impaired dicing activity, a step upstream in the RNA silencing pathway that is normally associated to DRB proteins.

**Figure 7:**
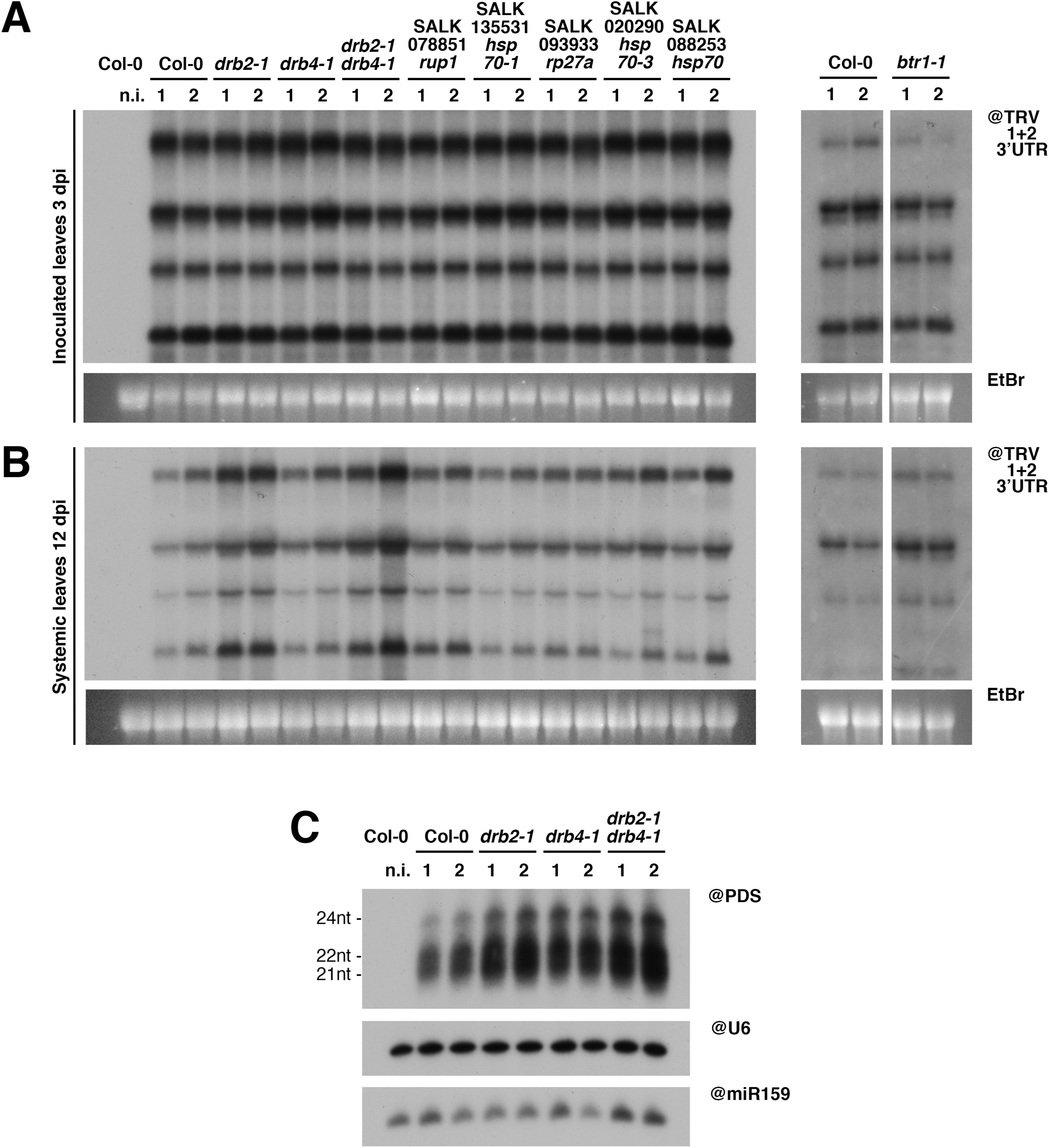
Knock-out of DRB2 causes increased systemic accumulation of TRV in Arabidopsis, through a mechanism independent from small RNA biogenesis. Northern blot analysis of RNA from inoculated leaves **(A)** and systemically infected leaves **(B)** of Arabidopsis knock-out lines infected with TRV-PDS, 3 and 12 days post-infection (dpi), respectively. Previously published mutants are indicated with their current name, while the others are indicated with their SALK nomenclature. Each sample is a pool of 4-5 plants, and two samples were analyzed per genotype (1 and 2), per time point. EtBr staining was used as loading control. **(C)** PAGE northern blot analysis of small RNA from the corresponding samples in (B). snU6 and miR159 were used as loading controls. N.i.: non-infected. Source data is available with the Blotting Source Data.

### DRB2 over-expression drastically reduces the accumulation of various plant RNA viruses

The absence DRB2 resulting in increased TRV accumulation (**Figure 7**), we next tested whether AtDRB2 over-expression could negatively impact infection by TRV and possibly by other distantly-related RNA viruses. To this end, agro-infiltrated *N. benthamiana* leaves transiently expressing DRB2:tRFP or tRFP were mechanically inoculated with the viruses of interest, and three days after infection leaf disks were collected. Northern blot analysis revealed that in tissues infected by TRV-PDS (**Figure 8A**), tomato bushy stunt virus (TBSV, **Figure 8B**), potato virus X (PVX, **Figure 8C**) and grapevine fanleaf virus (GFLV, **Figure 8D**), over-expression of DRB2:tRFP lead to a dramatic decrease in virus accumulation compared to over-expression of tRFP alone. This effect was particularly prominent for TBSV, PVX and GFLV, despite the presence of comparable amounts of DRB2:tRFP (**Figure 8E**). Remarkably, confocal microscopy of B2:GFP-expressing *N. benthamiana* leaves transiently over-expressing DRB2:tRFP and infected with TBSV showed that DRB2:tRFP co-localizes with VRCs (**Figure 8F**) that are structurally very different from those produced upon TRV infection (**Figure 4B)**. To confirm that these are indeed TBSV VRCs, which are known to form on peroxisome membranes [12], we generated a clone to express a tRFP-SKL peroxisome marker [48]. Expression of this marker in B2:GFP-expressing *N. benthamiana* leaves subsequently infected with TBSV reveal that B2-labeled VRCs are indeed localized on the surface of peroxisomes, that in infected conditions appear to group into large multi-peroxisome clusters (**Figure 8G**). These results clearly show that *At*DRB2 localizes to VRCs from different viruses and is a broad-ranged and potent antiviral effector.

**Figure 8:**
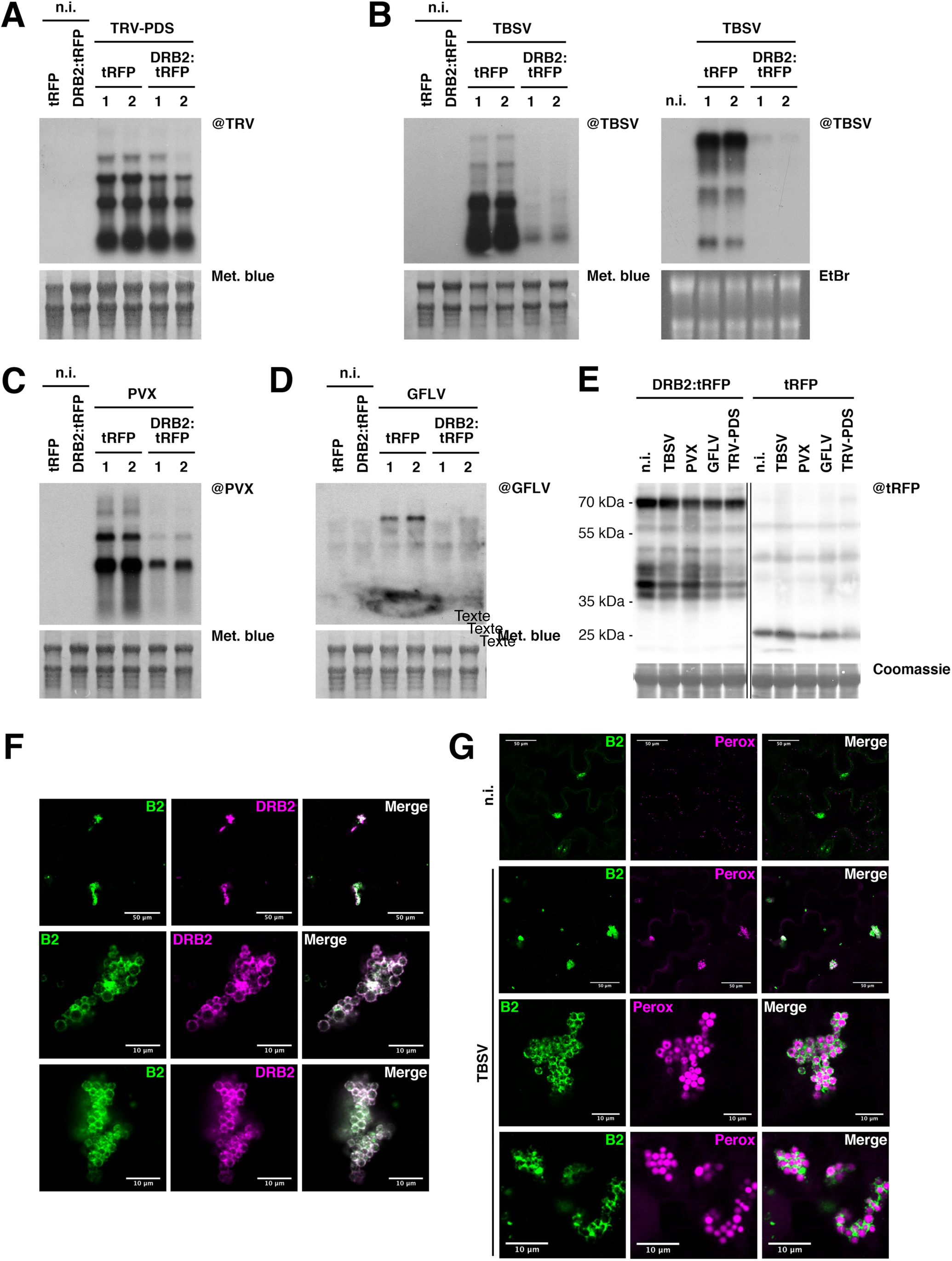
Over-expression of DRB2 in *N. benthamiana* leaves drastically reduces accumulation of a wide range of RNA viruses. **(A)** Northern blot analysis of RNA from *N. benthamiana* leaf disks 4 days after transient transformation with *35S:tRFP* or *35S:DRB2:tRFP* and 3 dpi with TRV-PDS (except for n.i.: non-infected). Each sample is a pool of 40-50 leaf disks from 4-5 leaves. In the case of the virus-infected leaves, two samples were analyzed per condition (indicated with 1 and 2). Methylene blue staining of the membrane was used as loading control. **(B)** As in (A), but after rub-inoculation with tomato bushy stunt virus (TBSV). Two independent biological replicates are shown with either methylene blue or EtBr staining of the membranes as loading control. **(C)** As in (A), but after rub-inoculation with potato virus X (PVX). **(D)** As in (A), but after rub-inoculation with grapevine fanleaf virus (GFLV). **(E)** Western blot analysis on protein extracts from the samples analyzed in (A-D), to detect tRFP. Coomassie blue staining was used as loading control. Source data is available with the Blotting Source Data. (**F**) Laser confocal microscopy acquisitions of B2-labeled TBSV replication complexes, from 35S:*B2:GFP*/*N. benthamiana* plants transiently expressing DRB2:tRFP and infected with TBSV. Scale bars indicate 50 (top) or 10 μm (middle and bottom). **(G)** As in (F), but from plants (non-infected in the top row, TBSV-infected in the rest) transiently expressing the peroxisome marker tRFP-SKL. Scale bars indicate 50 (top two acquisitions) or 10 μm (bottom two acquisitions). Additional acquisitions can be found with the Microscopy Source Data.

## DISCUSSION

We have here provided a description of a novel approach toward the identification of VRC-associated proteins through the isolation of replicating viral dsRNA during genuine infection, and validated the localization of most of the candidates through a rapid, robust and simple system. We also showed that one of the proteins we identified as associated to viral dsRNA, DRB2, has antiviral activity against several RNA viruses that belong to different families including *Secoviridae* (GFLV), *Virgaviridae* (TRV), *Tombusviridae* (TBSV) and *Alphaflexiviridae* (PVX). Although the proof of concept for our approach to identify VRC-associated proteins is established here only for TRV, it should be compatible with any plant virus as long as it is able to produce dsRNA during its replication cycle. Importantly, it does not involve as a prerequisite any modification of viral genomes, the production of infectious clones or the specific tagging of viral protein. Also, considering that the isolation of viral dsRNA and associated proteins is achieved indirectly by anti-GFP antibodies, there is no requirement for virus- or dsRNA-specific antibodies in the process. Hopefully this experimental approach will provide future investigators with a universal tool to successfully explore the proteome associated to the replication complexes of their favorite RNA virus, which can then be studied more in detail to discover the function of VRC-associated proteins and their involvement in the viral life cycle. As hosts, 35S:*B2:GFP*/*A. thaliana* (this study) and 35S:*B2:GFP*/*N. benthamiana* [11] are compatible with numerous plant virus species. If needed, the systems could be easily adapted to other plant species, as long as they accommodate stable transformation.

We have shown that ectopic expression of B2:GFP greatly increases the accumulation of TRV RNA both in *A. thaliana* and *N. benthamiana*. Given the activity of the 73 amino-acid double-stranded binding domain of B2 as a VSR (this work), it is tempting to ascribe TRV over-accumulation simply as a consequence of RNA silencing suppression and subsequent enhanced viral replication. While this is very probably the case, it cannot be excluded that B2:GFP increases TRV accumulation by RNAi-independent means, such as stabilization of dsRNA or its protection from other host defensive pathways [49]. Preliminary data also suggest that enhanced viral replication may not be restricted to TRV but likely applies to other RNA virus (Ritzenthaler, unpublished).

The drastic effect of B2:GFP on TRV infection can be viewed as a double-edged sword in relation to its use as bait to pull down VRCs. On one hand, this effect may introduce biases of both quantitative and qualitative nature, such as the unspecific association to VRCs of host proteins that do not play a role during infection in wild-type conditions or changes in the accessibility or protein complement of replicating RNA, for example. On the other hand, the over-accumulation of TRV dsRNA constitutes a real advantage for the study of VRCs. In fact, increased viral replication is in favor of (i) a better immunoprecipitation efficiency, (ii) an enhanced detection of protein partners by mass spectrometry and (iii) an improved visualization of VRCs with test candidates. While these biases clearly need to be taken into account, we strongly believe that overall this approach has far more advantages than drawbacks.

The abundance of TRV replicase detected in the IPs (624 reads, Figure 2A) is, in our opinion, confirmation of the robustness of the experiment in terms of VRC yield and integrity. The abundant detection of the coat protein suggests that it either plays a direct role in TRV replication or that the VRCs present in the IPs contain not only full-length dsRNA, but also (+)ssRNA that is being encapsidated, possibly during or just after separation from the (-) strand. However, despite the use of detergent during the IPs, it is possible that we have pulled down proteins present on membranes or complexes close to the replication organelles but not actually part of them.

Remarkably, among the nine candidate *A. thaliana* proteins detected following B2 IP and tested in this work, only one, HSP70, failed to accumulate in VRCs despite a high spectral count. HSP70 has been linked to viral infection in a number of studies [5, 42] and found to directly bind the viral replicase of at least two viruses [50, 51]. It is possible that the tRFP could have disrupted the function of the protein. Another possibility is that the *A. thaliana* HSP70 (AT3G12580) may not be fully functional when expressed in the heterologous host *N. benthamiana*.

All remaining 8 candidates were specifically redistributed upon infection, suggesting involvement of these factors in the viral life cycle. Their localization patterns can be divided into three groups: perfect co-localization (DRB2 and DRB4), partial co-localization (HSP70-1, HSP70-3, BTR1, RUP1 and RP27a) and proximity (NFD2).

Perfect co-localization most likely reflects the direct association of DRB2 and DRB4 on replicating dsRNA within the VRCs. This result is in line with the experimentally verified ability of DRB2 and DRB4 to bind dsRNA [35, 52] and of DRB4 to bind TYMV dsRNA *in vivo* [40]. DRB2 was recently shown in *A. thaliana* to re-localize to cytoplasmic punctate bodies upon infection by TuMV, TSWV and TYMV [8] and DRB4 to re-localize from nuclei to cytoplasmic VRCs upon TYMV infection [40]. While DRB4 plays a role in antiviral defense [40, 41] as part of the RNA silencing machinery, the function of DRB2 recruitment to replication complexes remains to be uncovered. Although additional experiments are required to confirm the direct association of DRB2 and DRB4 to TRV replicating dsRNA and DRB2 to TBSV replicating dsRNA, our data suggest that host proteins including antiviral defense protein such as DRBs may have access to viral dsRNA within replication organelles including TBSV-induced spherules. This potentially questions the suggested function (or at least efficiency) of replication organelles as protective structures against degradation by cellular RNases and detection by putative dsRNA sensors that trigger antiviral responses [12, 53, 54]. It is conceivable that B2:GFP and the DRB proteins gain access to viral dsRNA at early stage of replication organelle morphogenesis before replication complexes become eventually fully protected.

While the precise molecular pathways linking DRB2 to VRCs remain to be uncovered, we have shown through genetic ablation and over-expression that this protein is a key element in the host’s restriction of viral systemic infection. We have also shown that the antiviral activity of DRB2 likely does not involve Dicer function, since viral siRNA production remains unchanged upon knock-out of DRB2. Our data, however, does not rule out a possible involvement of DRB2 in steps of the RNA interference pathway that are downstream of Dicer processing. Whatever the molecular mode of action of DRB2, our over-expression experiments have shown that heightened production of this protein *in planta* drastically reduces the accumulation of viruses belonging to various families. Therefore, we believe that further study of DRB2 and its use as a biotech tool in crop defense against viral infection hold substantial potential.

The pattern of partial co-localization, observed for HSP70-1, HSP70-3, BTR1, RUP1 and RP27a/Ubiquitin, consisted in the localization at the B2-labeled VRCs *per se*, as well as features in close proximity, generally designated as “viral factories”. In the case of PVX, the dsRNA-containing replication complexes reside within larger viral factories harboring other viral proteins and viral ssRNA [11]. In general, these viral factories are most likely the hub for a plethora of viral activities beyond RNA replication *sensu stricto,* such as translation, encapsidation, etc…, and which likely require specific host-encoded proteins. Our work suggests that HSP70-1, HSP70-3, BTR1, RUP1 and RP27a/Ubiquitin may play such functions during replication of TRV and possibly other viruses. Indeed, HSP70-3 and BTR1 have been shown to interact with TuMV replicase [16] and ToMV ssRNA [30], respectively. Similarly, ubiquitin and the ubiquitin pathway have been shown in a number of studies to play important roles in plant virus life cycle, both pro-viral and anti-viral, the details of which are exhaustively reviewed in [55, 56]. Finally concerning NFD2, the pattern of proximity suggests that this protein may indirectly be involved in viral factory function without directed association with viral dsRNA per se. The fact that genetic knock-out of most of these factors did not lead to drastic changes in viral RNA accumulation (with the notable exception of DRB2) does not rule out their involvement in viral functions despite their localization to VRCs. They could act redundantly with other proteins, or could affect parameters that do not perturb viral RNA accumulation, to name but a few possibilities. The genetic dissection of the roles played by the proteins here identified, through experiments including IP and mass spectrometry of tagged alleles of these factors in different genetic backgrounds, is outside the scope of this manuscript and will be addressed in further studies.

## MATERIALS AND METHODS

### Golden Gate pEAQΔP19 vector construction

Binary vector pEAQΔP19-GG was obtained by, (i) removing 3 SapI restriction sites present in pEAQ-*HT* [57], (ii) inserting a Golden Gate cassette (similar to Gateway without AttR1/2) with SapI sites at extremities and (iii) removing P19. Two silent substitutions into Neomycin phosphotransferase (nptII) gene and one substitution near the origin of replication (ColE1) were produced by PCR mutagenesis using Phusion polymerase in GC buffer supplemented with 5% DMSO (primers in **Supplementary Table 3**, n°595-596 and 638-641) in order to obtain plasmid pEAQ-HT-ΔSapI. A Golden Gate cassette amplicon (pEAQ-HT as matrix, see primer n°589+642 **Supplementary Table 3**) was inserted via AgeI/XhoI restriction sites in pEAQ-HT-ΔSapI. Finally, P19 was excised by double restriction EcoNI/SgsI (FD1304, FD1894, Thermo Scientific), extremities were filled in with Klenow fragment (EP0051, Thermo Scientific), supplemented with dNTPs, followed by a ligation step and transformation in *E. coli* (ccdB Survival strain, Invitrogen).

### Plant material

Transgenic 35S:*B2:GFP*/*N. benthamiana* plants were previously described [11]. Transgenic *A. thaliana* plants (Col-0 line and genetic backgrounds including mutants of the core antiviral Dicer-Like genes, *dcl2-1*, *dcl4-2* and triple *dcl2-1/dcl3-1/dcl4-2* lines - [29] expressing 35S:*B2:GFP* were generated using same plasmid (pEAQΔP19-B2:GFP) and agrobacteria described in [11], following floral dip transformation [58] with addition of PPM (Plant Preservative Mixture, Plant Cell Technology) at 2 ml/L in MS medium. Individual Arabidopsis transformed lines were self-pollinated to generate (F3) plants homozygous for the transgene.

### Cloning of candidate genes

Candidate genes were amplified from *A. thaliana* genomic DNA with primers designed to contain SapI restriction sites compatible with Golden Gate cloning [59] and adapters necessary for ligation to an N-terminal or C-terminal tag (primer list in **Supplementary Table 3**). In the case of genes containing SapI restriction sites, silent mutations were introduced to remove these sites through overlap PCR. In parallel, tRFP was amplified with primers designed to contain SapI restriction sites, adapters for ligation to the N- or C-terminal end of the candidate gene and a peptide linker (GGGSGGG amino acid sequence) between tRFP and the candidate gene. tRFP-SKL was generated by adding the bases to encode the SKL tripeptide in the reverse primer before the stop codon. PCR products were purified from agarose gel and used in a Golden Gate reaction containing the candidate gene, tRFP, binary vector pEAQΔP19-GG, SapI (R0569L, New England Biolabs), CutSmart buffer (New England Biolabs), T4 DNA ligase 5U/µl (EL0011, Thermo Scientific) and 0.5 or 1 mM ATP (R0441, Thermo Scientific). Golden Gate reaction cycling: 10 cycles of 37°C 10 min, 18°C 10 min; 18°C 50 min, 50°C 10 min, 80°C 10 min. Following transformation in *E. coli* (TOP10 strain, Invitrogen), purification and sequencing, plasmids were transformed into *A. tumefaciens* strain GV3101. pEAQΔP19-B2:RFP served as matrix to generate plasmid pEAQΔ P19-B2mut:RFP as described in [11](primers n°631-632, **Supplementary Table 3**).

### Plant inoculation and infection

For fluorescence microscopy experiments, leaves of 5-6 week-old 35S:*B2:GFP/N. benthamiana* were infiltrated with *A. tumefaciens* GV3101 carrying plasmid pEAQΔP19 containing the tagged gene of interest, at absorbance_600nm_ (A_600_) of 0.2. Prior to inoculation, bacteria were incubated in 10 mM MES pH 5.6, 10 mM MgCl_2_, 200 μM acetosyringone for 1 hour. TRV infection was initiated upon agro-infection with bacteria carrying plasmids expressing the two viral genomic RNAs [28], at A_600_ 0.01 each. This method of virus delivery was chosen because it results in homogenous and ubiquitous infection throughout the inoculated tissue. 3-4 days post-inoculation, leaf disks were collected and observed by confocal microscopy. *A. thaliana* infection was carried out as for *N. benthamiana*, with the difference that *A. tumefaciens* was induced by incubating 5-6 hours in induction medium (10.5 g/L K_2_HPO_4_, 4.5 g/L KH_2_PO_4_, 1 g/L (NH_4_)_2_SO_4_, 0.5 g/L sodium citrate, 0.1 g/L MgSO_4_, 0.4% glycerol, 0.1 g/L MES, 200 μM acetosyringone), and bacteria used at A_600_ 0.5 each. Systemically infected leaves were harvested 12-13 dpi.

For the experiments shown in Figure 8, virus infection was carried out by rub inoculation: the day following agro-infiltration with 35S:*tRFP,* 35S:*tRFP-SKL* or 35S:*DRB2:tRFP*, the abaxial side of the infiltrated leaves was mechanically inoculated. The inoculum was obtained by grinding frozen *N. benthamiana* tissues infected with TBSV, TRV-PDS, PVX-GFP or GFLV in 50 mM sodium phosphate buffered at pH 7 (except for TBSV, at pH 5.8). *N. benthamiana* plants were kept in a greenhouse at 22-18°C, 16h/8h light/dark photoperiod, while *A. thaliana* were kept in a neon-lit growth chamber at 22-18°C, 12h/12h light/dark photoperiod.

### Immunoprecipitation

Immunoprecipitations were performed as previously described [21], with minor modifications. 0.15 g of young rosette leaves were ground in liquid nitrogen, homogenized in a chilled mortar with 1 ml lysis buffer (50 mM Tris–HCl, pH 8, 50 mM NaCl, 1% Triton X-100) containing 1 tablet/50 ml of protease inhibitor cocktail (Roche), transferred to a tube and incubated for 15 min at 4°C on a wheel. Cell debris was removed by two successive centrifugations at 12000*g* for 10 min at 4°C, after which an aliquot of supernatant was taken as input fraction. The remaining extract was incubated with magnetic microbeads coated with monoclonal anti-GFP antibodies (μMACS purification system, Miltenyi Biotech, catalog number #130-091-125) at 4°C for 20 min. Sample was then passed through M column (MACS purification system, Miltenyi Biotech) and an aliquot of the flow-through fraction was taken. The M column was then washed 2 times with 500 μl of lysis buffer and 1 time with 100 μl of washing buffer (20 mM Tris–HCl, pH 7.5). The beads and associated immune complexes were recovered by removing the M column from the magnetic stand and passing 1 ml Tri Reagent (for subsequent RNA analysis – see dedicated section) or 200 μl hot 1X Laemmli buffer (for protein analysis – see dedicated section). 4X Laemmli buffer was added to input and flow-through fractions before protein denaturation for 5 min at 95°C.

### RNA extraction and analysis

RNA from total and immunoprecipitated fractions was performed with Tri-Reagent (Sigma) according to manufacturer’s instructions. Briefly, 0.2 g tissue were ground in liquid nitrogen and homogenized in 1 ml Tri-Reagent, 400 μl of chloroform were added, and sample was thoroughly shaken for 2 min. After 10 min spin at 13000 rpm, 4°C, supernatant was added to at least 1 vol isopropanol (and 1.5 μl glycogen in the case of immunoprecipitated samples - IP) and incubated 1 hour on ice (O/N for IP). After 15 min spin at 13000 rpm, 4°C (30 min for IP), pellet was washed in 80% ethanol, dried and resuspended in water. RNA was analyzed by northern blot (denaturing agarose gel to detect high molecular weight RNA, denaturing PAGE to detect low molecular weight RNA) and northwestern blot (native agarose gel to detect long double-stranded RNA). In northern blot, miRNA were detected through DNA oligonucleotides labeled with γ-^32^P-ATP using T4 PNK (see **Supplementary Table 3**). TRV genomic and subgenomic RNAs were detected in the same way, with an oligonucleotide complementary to a part of the 3’UTR sequence common to RNA1 and RNA2. The same was done for TBSV. TRV-PDS-derived siRNA were detected through PCR-amplified *A. thaliana* PDS sequence labeled by random priming reactions in the presence of α-^32^P-dCTP. The same was done to detect PVX and GFLV RNA in Northern blot. In northwestern blot, dsRNA were detected through recombinant Strep-Tagged FHV B2, as previously described [11].

### Protein extraction and analysis

Proteins from total fractions were extracted as previously described [60]. Immunoprecipitated proteins for mass spectrometry analysis were isolated as described above, then denatured 5 min at 95°C. Immunoprecipitated proteins from RNA IP were obtained by collecting the phenolic phase following Tri-reagent/chloroform extraction, adding 3-4 vol acetone and incubating at −20°C O/N. After centrifugation (13000 rpm, 15 min, 4°C) pellet was washed in 80% acetone and resuspended in 1X Laemmli. Proteins were resolved by SDS-PAGE and electro-blotted onto Immobilion-P membrane. This was incubated with the appropriate antibodies (anti-GFP polyclonal antibody and anti-tRFP antibody, Evrogen, reference # AB233) and revealed with Roche LumiLight ECL kit after incubation with secondary antibody.

### Mass spectrometry analysis and data processing

Proteins were digested with sequencing-grade trypsin (Promega) and analyzed by nanoLC-MS/MS on a TripleTOF 5600 mass spectrometer (Sciex, USA) as described previously [61]. Data were searched against the TAIR v.10 database with a decoy strategy (27281 protein forward sequences). Peptides were identified with Mascot algorithm (version 2.5, Matrix Science, London, UK) and data were further imported into Proline v1.4 software (http://proline.profiproteomics.fr/). Proteins were validated on Mascot pretty rank equal to 1, and 1% FDR on both peptide spectrum matches (PSM score) and protein sets (Protein Set score). The total number of MS/MS fragmentation spectra was used to quantify each protein from at least three independent biological replicates. A statistical analysis based on spectral counts was performed using a homemade R package as described in [62]. The R package uses a negative binomial GLM model based on EdgeR [63] and calculates, for each identified protein, a fold-change, a p-value and an adjusted p-value corrected using Benjamini-Hochberg method.

### Confocal laser scanning microscopy

Observations of leaf disks were carried out using Zeiss LSM700 and LSM780 laser scanning confocal microscopes. eGFP was excited at 488 nm, while tRFP was excited at 561 nm. Image processing was performed using ImageJ/FIJI, while figure panels were assembled with Adobe Photoshop.

## DATA AVAILABILITY

All data are available in the manuscript and in Supplementary files. Raw data from gels and blots can be found with the Blotting Source Data file. Additional confocal acquisitions for each candidate tested can be found in the Microscopy Source Data file. Raw/unprocessed confocal images (.lsm format) and mass spectrometry data used to assemble all figures have been deposited in the public repository Zenodo. They can be accessed at the DOI: TO BE GENERATED.

## ACKNOWLEDGEMENTS

We thank David Gilmer for providing anti-GFP (Rocco) polyclonal antibodies. Special thanks to Hélène Zuber for the R-script that enabled the statistical analysis of co-IP experiments. We also would like to thank Philippe Hammann for assistance in immunoprecipitation experiments, and Jerome Mutterer and Mathieu Erhardt for assistance in laser confocal microscopy.

## FUNDING

Funding Disclosure: This work was co-funded by the CNRS and the European Regional Development Fund (ERDF) in the framework of the INTERREG IV and V Upper Rhine programs (BACCHUS and VITIFUTUR projects). B.M. benefited from an IdEx postdoctoral fellowship from the Université de Strasbourg. M.C. benefited from the European Research Council under the European Union’s Seventh Framework Programme (FP7/2007-2013) / ERC advanced grant to PG agreement n°[338904]. M.I. and M.C. also benefited from funds from LABEX: ANR-10-LABX-0036_NETRNA.

## COMPETING INTERESTS

The authors declare that the research was conducted in the absence of any commercial or financial relationships that could be construed as a potential conflict of interest.

## AUTHOR CONTRIBUTIONS

Study conception and design: M.I. and C.R.; generation of transgenic *A. thaliana* lines: B.M.; immunoprecipitation experiments: M.I.; RNA and protein extraction, northern and north-western blotting: M.I. and M.C.; western blotting: M.I., M.C. and H.S.; mass spectrometry: L.K.; statistical analysis of co-IP data: L.K. and H.S.; molecular cloning: M.I., B.M. and V.P.; recombinant B2 production: V.P.; *N. benthamiana* inoculation and infection: M.I. and M.C.; laser confocal microscopy: M.I.; data analysis: M.I., M.C., B.M., L.K., H.S., V.P., P.D. P.G., C.R. writing: M.I. and C.R.; supervision: C.R.; funding acquisition: C.R., P.G. and P.D.

---

**Supplementary Table 1.**

List of proteins detected by mass spectrometry in GFP pull-downs from 35S:*GFP*/Col-0 and 35S:*B2:GFP*/Col-0 plants infected with TRV-PDS. Here are shown only proteins present exclusively in B2:GFP or with a B2:GFP/GFP detection ratio ≥ 2. The candidates further investigated in this study are highlighted. A more detailed legend is present at the top of the spreadsheet.

**Supplementary Table 2.**

List of proteins detected by mass spectrometry in GFP pull-downs from 35S:*GFP*/Col-0 and 35S:*B2:GFP*/Col-0 plants in both non-infected and TRV-infected conditions. The IP from non-infected 35S:*GFP*/Col-0 vs 35S:*B2:GFP*/Col-0 plants was performed in a different experiment from that on TRV-infected plants (shown in Supplementary Table 1). The proteins detected have been sorted according to their presence/absence in the different genotypes and conditions. The candidates further investigated in this study are highlighted. A more detailed legend is present at the top of the spreadsheet.

**Supplementary Table 3.**

List of primers and probes used in this study.

**Supplementary Figure 1: (A)** Western blot analysis to detect GFP in protein extracts from 35S:*B2:GFP* transgenic *A. thaliana* lines in Col-0 and *dcl2-1*, *dcl4-2* and triple *dcl2-1/dcl3-1/dcl4-2* mutant backgrounds. Coomassie blue was used as loading control. **(B)** Northern analysis of low molecular weight RNA (PAGE gel) from the plants described in (A), to detect endogenous miRNA (miR159, miR160) and siRNA (TAS1, IR71). Probes were hybridized to the membrane through sequential stripping and probing. **(C)** Photos of the plants described in (A). **(D)** *N. benthamiana* leaf infiltrated with *A. tumefaciens* expressing GFP alone (top left) or in combination with P38, B2:tRFP or B2mut:tRFP, and illuminated with UV light. **(E)** Western blot analysis to detect GFP (top) and tRFP (bottom) in protein extracts from the infiltrated patches described in (E). **(F)** Northern analysis of high molecular weight RNA (agarose gel) from TRV-PDS systemically infected Col-0 and *dcl* mutant backgrounds. Source data is available with the Blotting Source Data.

**Supplementary Figure 2:** Western analysis to detect GFP in protein extracts from the input, flow-through and anti-GFP immunoprecipitated fractions obtained from TRV-PDS-infected 35S:*GFP*/Col-0 and 35S:*B2:GFP*/Col-0 plants, performed in three technical replicates. Coomassie staining was used as loading control. The proteins from the immunoprecipitated fraction were further analyzed by mass spectrometry, and the results are shown in Table 1. Source data is available with the Blotting Source Data.

**Supplementary Figure 3:** Laser confocal microscopy (20x objective) on 35S:*B2:GFP/N. benthamiana* non-infected (left) and TRV-PDS-infected (right) leaf disks transiently expressing **(A)** 35S:*HSP70:tRFP,* **(B)** 35S:*HSP70-1:tRFP*, **(C)** 35S:*HSP70-3:tRFP*, **(D)** 35S:*BTR1:tRFP*. Scale bars indicate 50μm. Additional acquisitions can be found with the Microscopy Source Data in the Supplementary Information.

**Supplementary Figure 4:** Laser confocal microscopy (20x objective) on 35S:*B2:GFP/N. benthamiana* non-infected (left) and TRV-PDS-infected (right) leaf disks transiently expressing **(A)** 35S:*RUP1:tRFP,* **(B)** 35S:*tRFP:RP27a*, **(C)** 35S:*tRFP:NFD2*. Scale bars indicate 50 μm. Additional acquisitions can be found with the Microscopy Source Data in the Supplementary Information.

**Supplementary Figure 5: (A)** Sequence coverage obtained by MS on the RP27a protein (P59271). Four distinct tryptic peptides have been validated by Mascot algorithm at FDR<1% (bold residues). The total number of spectra matching on the 4 peptides are displayed as green bars with a color code according to the replicate. The Ubiquitin domain [1–76] is highlighted in green. Tryptic cleavage sites (K/R) are underlined. **(B)** Multiple sequence alignment between RP27a sequence and UBQ1 to UBQ14 *A.thaliana* sequences (UniProtKB). Output generated with the MUSCLE tool (https://www.ebi.ac.uk/Tools/msa/muscle). **(C)** MS/MS spectrum corresponding to the ubiquitinated peptide [43–54] LIFAGK(Ub)QLEDGR identified by Mascot algorithm on RP27a protein (Score = 55.68, m/z=487.60, 3+, RT=42.97min). The fragmentation pattern involving the y-and b-ions validated by Mascot is displayed on the upper left corner. The di-glycine motif is highlighted by the GL abbreviation above the K-48 residue, as well as the mass difference between y(6) and y(7) fragments.

