## Supplementary figures for "Immunocapture of dsRNA-bound proteins provides insight into tobacco rattle virus replication complexes and reveals Arabidopsis DRB2 to be a wide-spectrum antiviral effector"

### Supplementary Figure 1

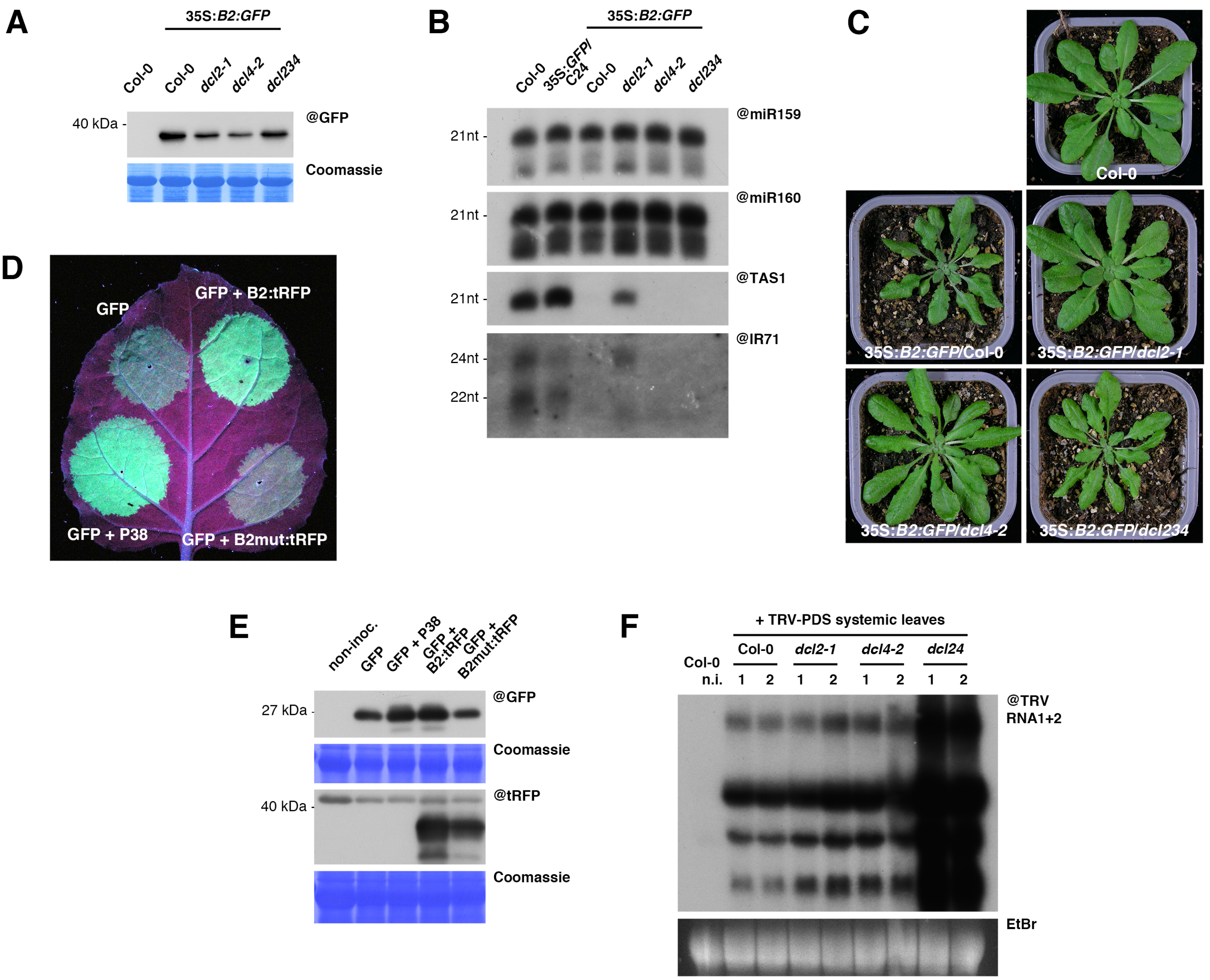

### Supplementary Figure 2

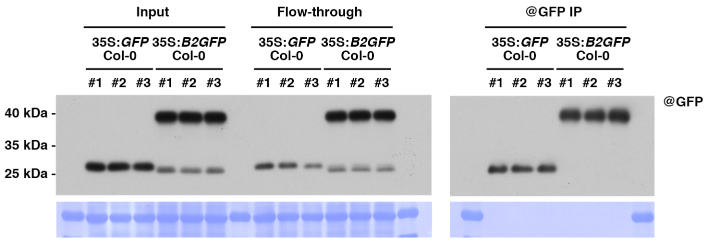

### Supplementary Figure 3

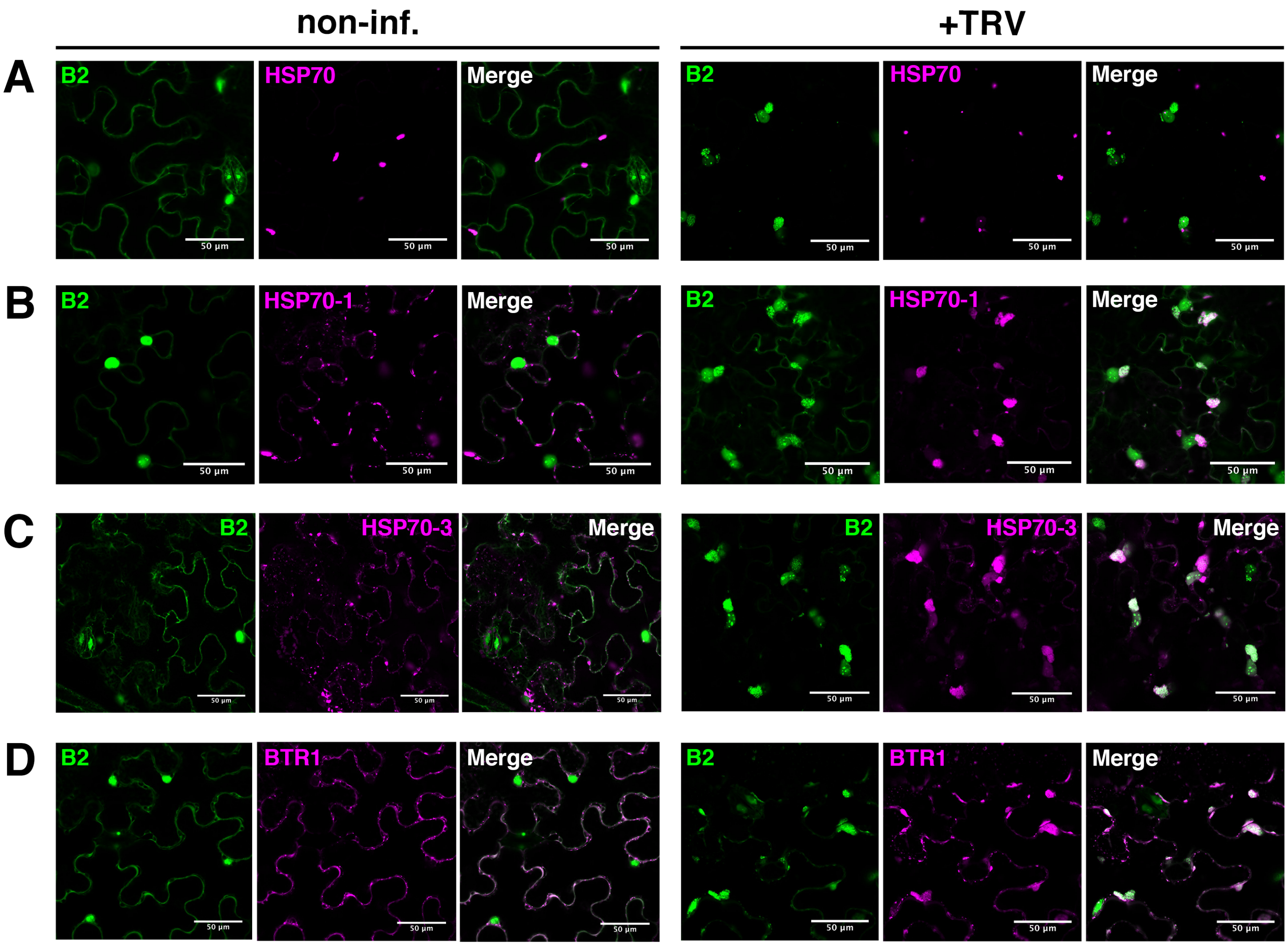

### Supplementary Figure 4

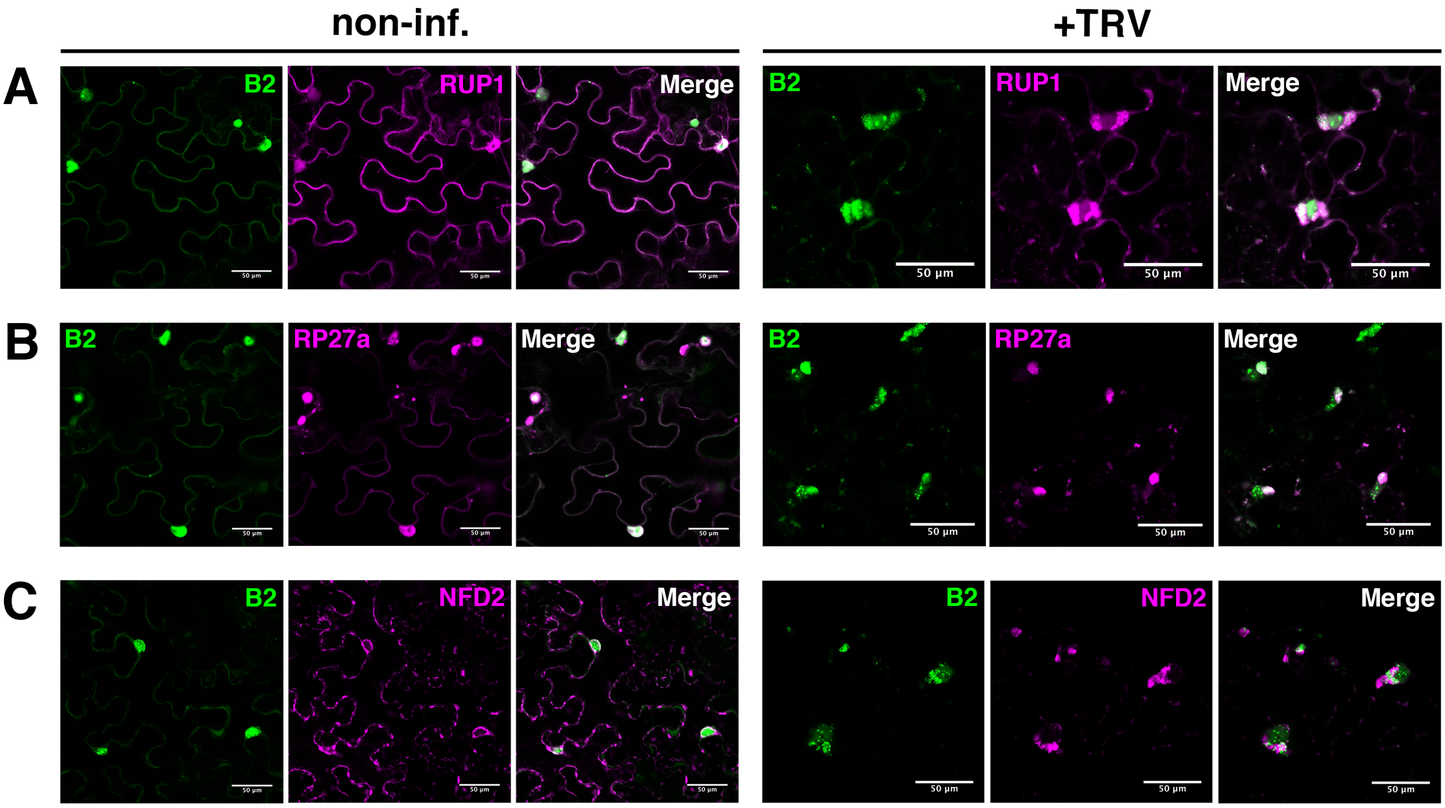

### Supplementary Figure 5

## A

>sp|P59271|R27AA_ARATH Ubiquitin-40S ribosomal protein S27a-1 (RPS27AA)

MQIFVKTLTGKTITLEVESSDTIDNVKAKIQDKEGIPPDQQRLIFAGKQLEDGRTLADYNIQKESTLHLVLR

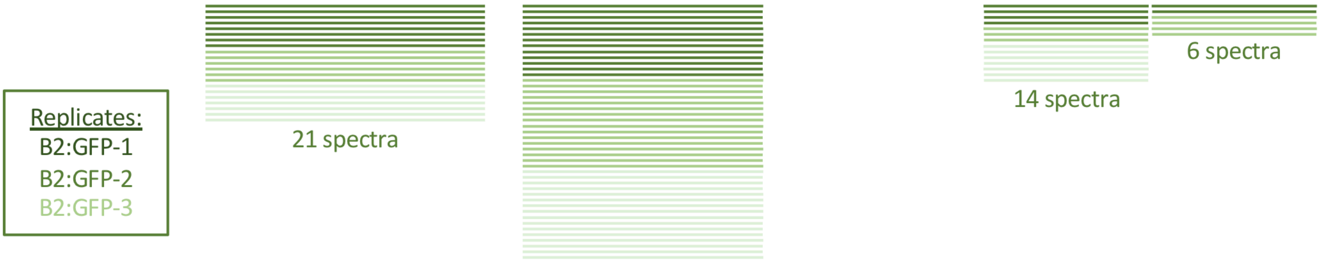


LRGGAKKRKKKTYTKPKKIKHTHKKVKLAVLQFYKVDGSGKVQRLKKECPSVSCGPGTFMASHFDRHYC
GKCGTTYVFKKADEE

## B

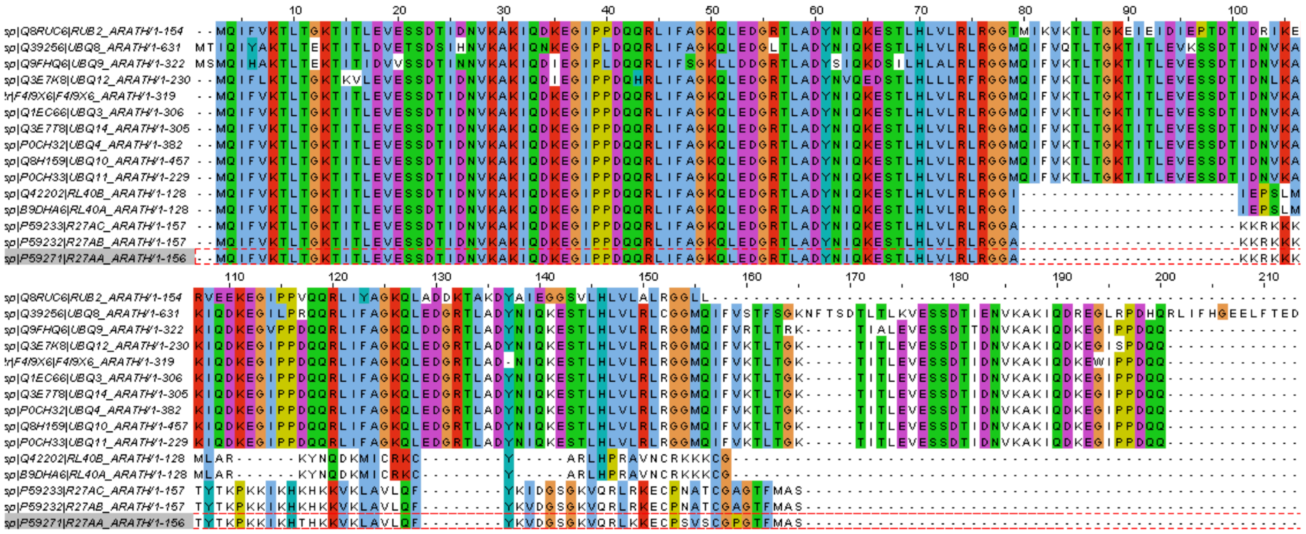

## C

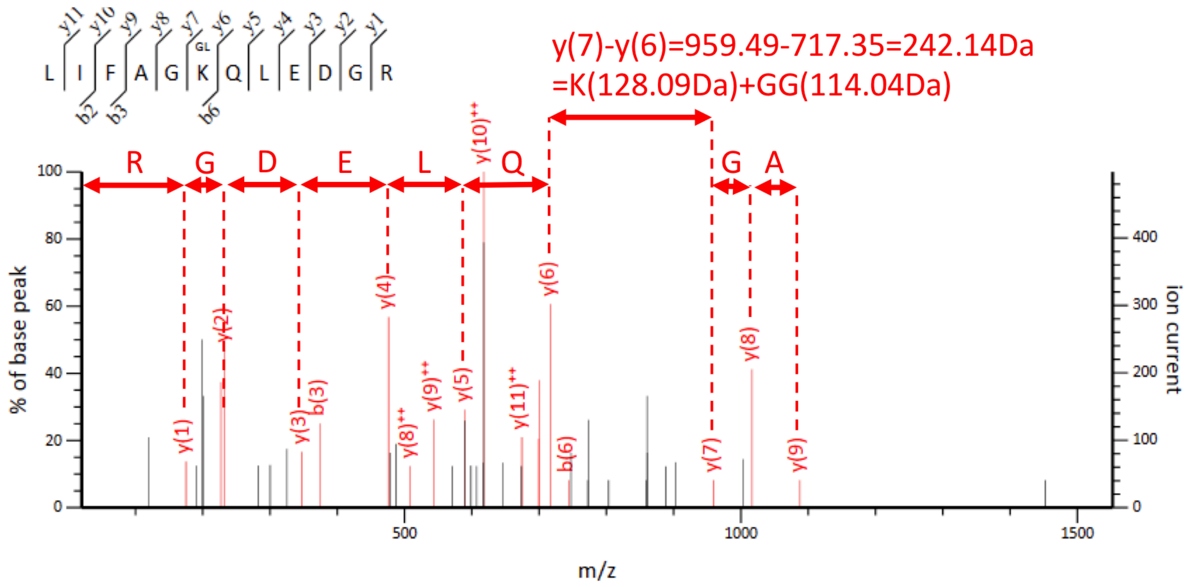
